# Reversal of cell, circuit and seizure phenotypes in a mouse model of *DNM1* epileptic encephalopathy

**DOI:** 10.1101/2023.04.14.536870

**Authors:** K. Bonnycastle, K.L. Dobson, E-M. Blumrich, A. Gajbhiye, E.C. Davenport, M. Pronot, M. Steinruecke, M. Trost, A. Gonzalez-Sulser, M.A. Cousin

## Abstract

Pathogenic heterozygous missense mutations in the *DNM1* gene result in a novel form of epileptic encephalopathy. *DNM1* encodes for the large GTPase dynamin-1, an enzyme with an obligatory role in the endocytosis of synaptic vesicles (SVs) at mammalian nerve terminals. Pathogenic *DNM1* mutations cluster within regions required for its essential GTPase activity, implicating disruption of this enzyme activity as being central to epileptic encephalopathy. We reveal that the most prevalent pathogenic mutation of *DNM1*, R237W, disrupts dynamin-1 enzyme activity and SV endocytosis when overexpressed in central neurons. To determine how this dominant-negative heterozygous mutant impacted cell, circuit and behaviour when expressed from its endogenous locus, we generated a mouse carrying the R237W mutation. Neurons isolated from heterozygous mice displayed dysfunctional SV endocytosis, which translated into altered excitatory neurotransmission and seizure-like phenotypes. Importantly, these phenotypes were corrected at the cell, circuit and *in vivo* level by the drug, BMS-204352, which accelerates SV endocytosis. This study therefore provides the first direct link between dysfunctional SV endocytosis and epilepsy, and importantly reveals that SV endocytosis is a viable therapeutic route for monogenic intractable epilepsies.

---

Heterozygous pathogenic missense mutations in the *DNM1* gene result in a specific form of developmental epileptic encephalopathy, characterised by severe to profound intellectual disability, hypotonia and epilepsy ^1–3^. Epilepsy in these individuals typically starts with infantile spasms progressing to Lennox-Gastaut syndrome. The *DNM1* gene encodes the large GTPase dynamin-1, a mechanochemical enzyme that undergoes a conformational change on GTP hydrolysis, providing force for the final stages of vesicle fission ^4, 5^. It has a modular structure with an N-terminal GTPase domain, followed by domains essential for self-assembly (middle and GTPase effector domains), membrane lipid binding (pleckstrin homology, PH) and protein interactions (C-terminal proline-rich domain). All domains perform key roles in mediating dynamin-1 function ^6–9^, however its GTPase activity is essential for SV fission reaction during endocytosis ^4, 5^. To date, all identified pathogenic mutations in the *DNM1* gene are localised to either the GTPase or middle domain, with one exception in the PH domain ^10^. Importantly, almost all individuals with these mutations in *DNM1* have intractable epilepsy ^1^, making the identification of novel therapeutic interventions an urgent unmet challenge.

Considering the essential role for dynamin-1 in SV endocytosis and the clustering of mutations within the GTPase domain in individuals with epileptic encephalopathy, a logical prediction is that defects in SV endocytosis underpin this disorder. However, to date, no determination of the role of these predicted dominant negative GTPase domain mutations has been performed at the neuron, circuit or behavioural level. To investigate this, we generated a mouse carrying the most prevalent mutation in the *DNM1* gene (R237W). Heterozygous *Dnm1*^+/R237W^ mice displayed defective SV endocytosis, altered neurotransmission and seizure-like activity. Critically, these phenotypes were all reversed by the small molecule BMS-204352. This novel preclinical model and potential therapy should provide impetus to future small molecule screening studies and clinical trials to generate novel interventions for *DNM1* epileptic encephalopathy.

## Results

### Dynamin-1 with a R237W mutation displays reduced basal GTPase activity

Heterozygous mutations in the *DNM1* gene give rise to a specific form of epileptic encephalopathy ^1^, however little is known regarding how these mutations translate into this neurodevelopmental disorder. To determine this, we investigated the most prominent pathogenic *DNM1* mutation in human disease - R237W ^1, 2^. The R237 residue is critical for GTPase function, directly participating in GTP binding and stabilising the transition state of this region during GTP hydrolysis ^11^. Because of this key role, we reasoned that substitution of a larger residue such as R237W would disrupt GTPase activity. To test this, we assayed the GTPase activity of full-length dynamin-1 that had been expressed in a heterologous expression system. This version of dynamin-1 was fused to the fluorescent protein mCerulean (Dyn1-mCer) and then immunoprecipitated using antibodies against the mCer moiety. As a positive control, we assessed the ability of the K44A dynamin-1 mutant (which is deficient in both GTP binding and hydrolysis ^12^) to hydrolyse GTP. When these experiments were performed, a level of baseline GTPase activity was discovered in immunoprecipitates of the mCer protein. However, this was increased two-fold in Dyn1_WT_-mCer immunoprecipitates (**Figure 1a,b**). In contrast, the Dyn1_K44A_-mCer mutant displayed a significantly reduced ability to hydrolyse GTP (**Figure 1a**). Importantly, Dyn1_R237W_-mCer displayed reduced GTPase activity at a similar level to Dyn1_K44A_-mCer (**Figure 1b**). Therefore, the R237W mutation disrupts the GTPase activity of dynamin-1.

**Figure 1.**
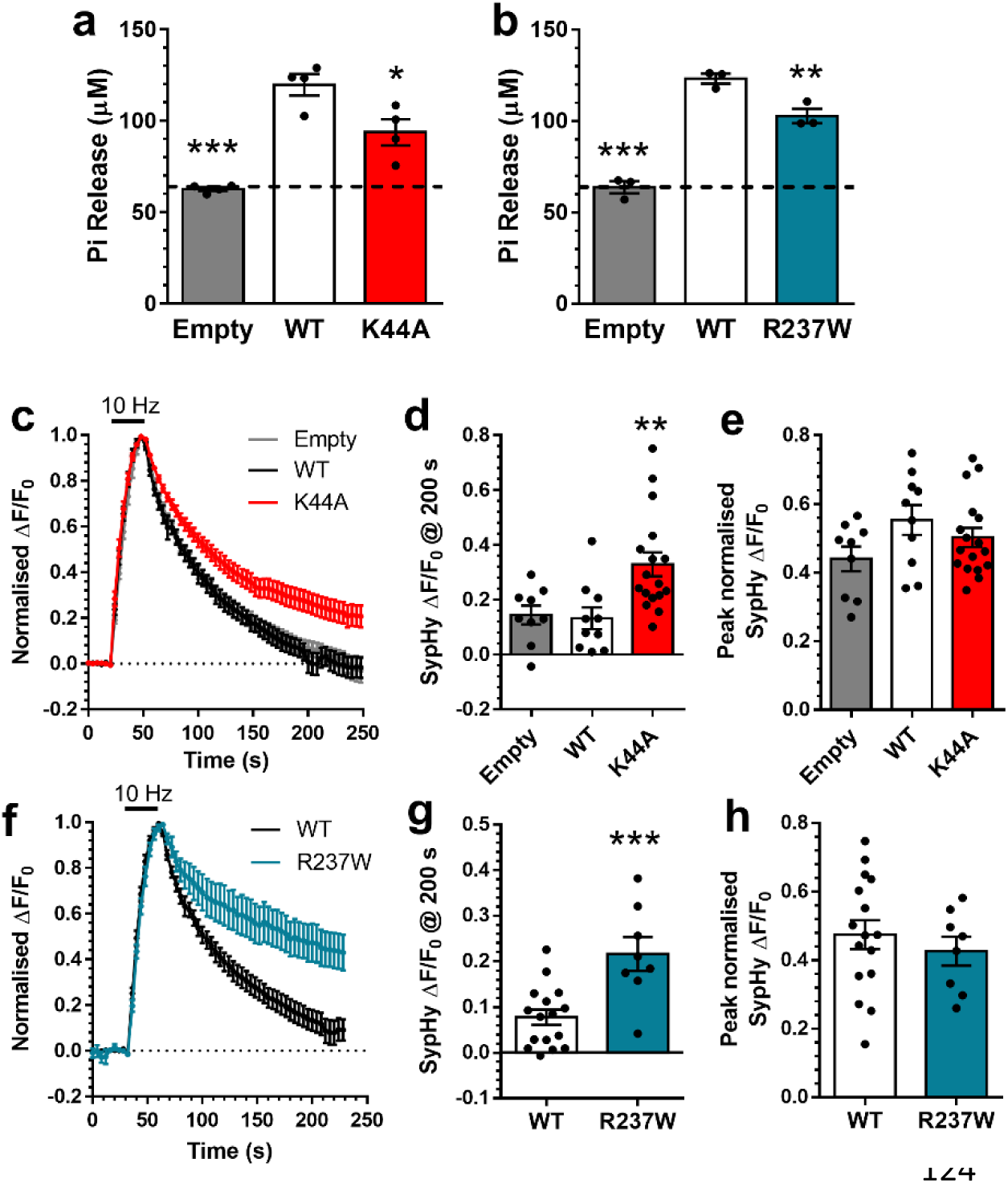
– *The R237W DNM1 GTPase domain mutation impairs SV endocytosis in a dominant negative manner.* (**a,b**) HEK293T cells were transfected with either mCer (Empty), Dyn1_WT_-mCer, Dyn1_K44A_-mCer or Dyn1_R237W_-mCer. After 48 h the cells were lysed and mCer was immunoprecipitated. The GTPase activity of the immunoprecipitate is displayed as released Pi ± SEM (one-way ANOVA, **a** all n=4 separate experiments, ***p<0.0001 WT to Empty, *p=0.0138 WT to K44A; **b** all n=3, ***p<0.0001 WT to Empty, **p=0.0093 WT to R237W). (**c-h**) Primary cultures of hippocampal neurons were transfected with synaptophysin-pHluorin (sypHy) and either mCer (Empty), Dyn1_WT_-mCer, Dyn1_K44A_-mCer or Dyn1_R237W_-mCer between 11-13 DIV. At 13-15 DIV, cultures were stimulated with a train of 300 action potentials (10 Hz). Cultures were pulsed with NH_4_Cl imaging buffer 180 s after stimulation. (**c,f**) Average sypHy response (ΔF/F_0_ ± SEM) normalised to the stimulation peak. Bar indicates stimulation. (**d,g**) The average level of sypHy fluorescence (ΔF/F_0_ ± SEM) at 200 s (**d** one-way ANOVA, n=9 Empty, n=10 Dyn1_WT_-mCer, n=17 Dyn1_K44A_-mCer, **p=0.0046 WT to K44A; **g** Unpaired t test, n=16 Dyn1_WT_-mCer, n=8 Dyn1_K44A_-mCer, *p=0.0011). (**e,h**) The peak level of sypHy fluorescence (ΔF/F_0_ ± SEM) normalised to the NH_4_Cl challenge (**e** one-way ANOVA, n=9 Empty, n=10 Dyn1_WT_-mCer, n=17 Dyn1_K44A_-mCer, all ns; **h** Unpaired t test, n=16 Dyn1_WT_-mCer, n=8 Dyn1_K44A_-mCer, p=0.48).

### Dynamin-1 with the R237W mutation inhibits SV endocytosis in a dominant-negative manner

Dynamin-1 GTPase activity is essential for SV endocytosis ^5^. To determine whether Dyn1_R237W_ exerts a dominant-negative effect on this process, and in an attempt to mimic heterozygous individuals with *DNM1* mutations, Dyn1-mCer mutants were overexpressed in primary cultures of wild-type hippocampal neurons, in approximately 2-3 fold in excess of endogenous dynamin-1 (**Extended Data Figure 1**). The genetically-encoded reporter synaptophysin-pHluorin (sypHy) was used to monitor activity-dependent SV recycling. SypHy is the SV protein synaptophysin that has an exquisitely pH-sensitive GFP inserted into a lumenal domain ^13^. The acidic interior of SVs result in the quenching of sypHy fluorescence in resting nerve terminals. During SV exocytosis, the reporter is exposed to the extracellular space and the subsequent unquenching provides a readout of SV fusion. SypHy remains fluorescent during endocytosis and is quenched on acidification of the newly formed SV. The speed of SV endocytosis is rate limiting when compared to SV acidification ^13, 14^, meaning that the loss of sypHy fluorescence is indicative of the rate of SV endocytosis.

Neurons were challenged with a train of 300 action potentials (10 Hz) with the total SV recycling pool revealed by subsequent application of an alkaline solution (NH_4_Cl). Neurons expressing the mCer empty vector displayed a typical sypHy response, with a rapid increase in fluorescence (reflecting SV exocytosis) followed by a slow decrease (SV endocytosis, **Figure 1c**). When neurons overexpressing Dyn1_WT_-mCer were monitored, there was no difference in either SV endocytosis (measured as the amount of sypHy left to retrieve (**Figure 1c,d**), or SV exocytosis (measured as the extent of the evoked sypHy peak as a proportion of the total SV pool, **Figure 1e**). In neurons expressing Dyn1_K44A_-mCer, SV endocytosis was significantly retarded, whereas SV exocytosis was unaffected (**Figure 1c–e**). When neurons overexpressing Dyn1_R237W_-mCer were assessed, SV endocytosis was greatly reduced compared to Dyn1_WT_-mCer control (**Figure 1f,g**). In contrast, there was no significant effect of Dyn1_R237W_-mCer on SV exocytosis (**Figure 1h**). Therefore, the pathogenic *DNM1* GTPase mutant R237W, has a selective, dominant-negative effect on SV endocytosis.

### *Dnm1*^+/R237W^ mice display defective SV endocytosis

The overexpression of Dyn1_R237W_-mCer in wild-type neurons does not accurately recapitulate the *in vivo* situation, where this mutation would be expressed via its endogenous locus. Furthermore, it does not allow a direct investigation of how reduced SV endocytosis could culminate in epileptic encephalopathy. To address this, we generated a mouse line that expressed an *Dnm1*^R237W^ allele, using CRISPR-Cas9 technology. Heterozygous *Dnm1*^+/R237W^ mice were fertile and were born in Mendelian proportions. No gross alterations in brain architecture were observed using Nissl staining (**Figure 2a**). Furthermore, the *Dnm1*^+/R237W^ mouse is not a hypomorph, since quantitative Western blotting and mass spectrometry analysis revealed no change in dynamin-1 expression in either primary hippocampal neurons from *Dnm1*^+/R237W^ mice, brain lysates from either 3 week- or 6 week-old *Dnm1*^+/R237W^ mice (**Extended Data Figure 2a,b**) or *Dnm1*^+/R237W^ nerve terminals (**Extended Data Table 1**).

**Figure 2.**
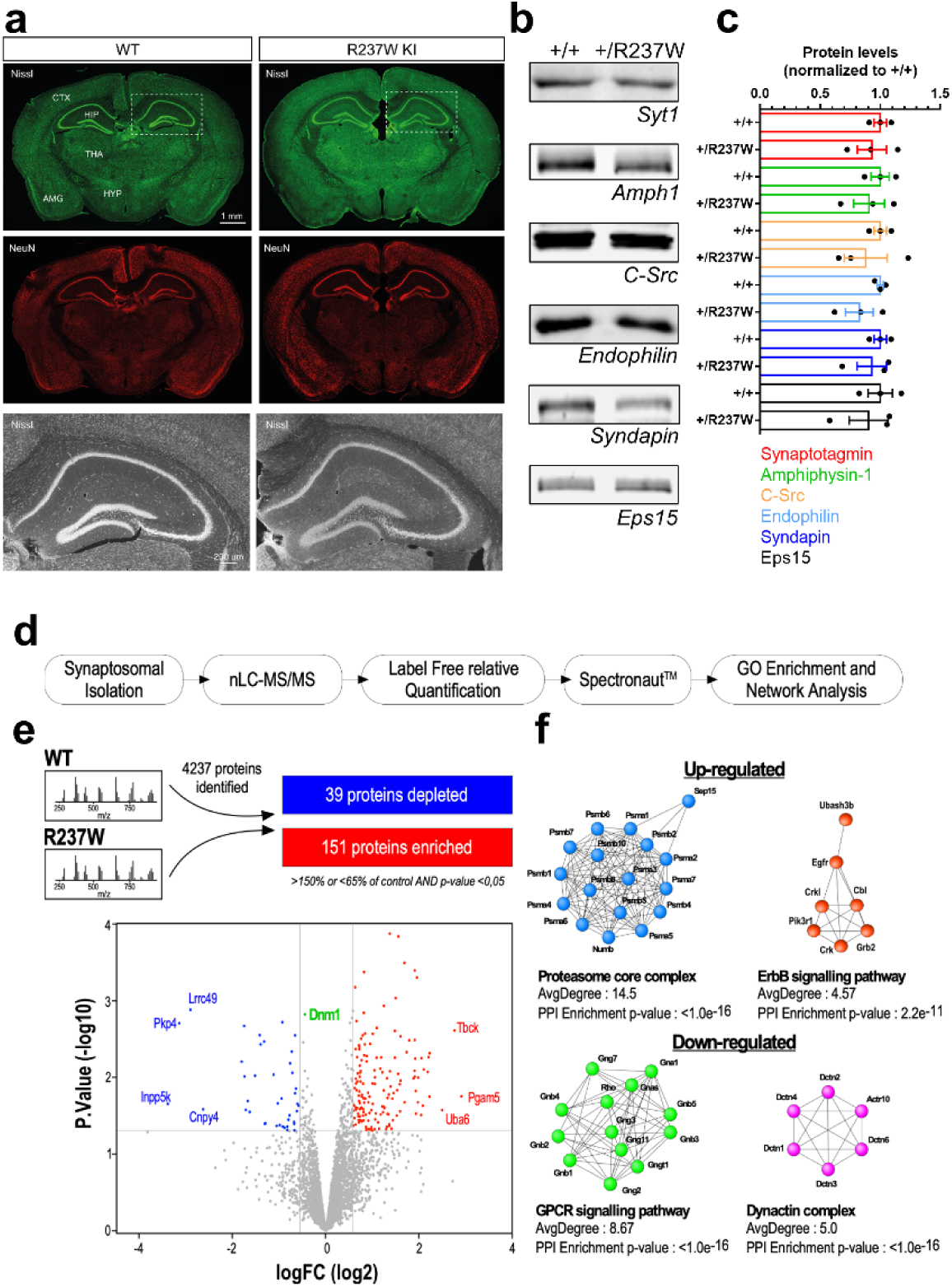
– *Dnm1*^+/R237W^ *mice display no gross abnormalities but altered protein expression.* (**a**) Brains from 2 month-old *Dnm1*^+/+^ (WT) and *Dnm1*^+/R237W^ mice were perfusion-fixed and 50 µm brain sections were stained with Nissl and a NeuN antibody to label gross neuronal architecture. Cortex (CTX), thalamus (THA), hypothalamus (HYP), amygdala (AMG) and hippocampus (HPC) are labelled, scale bar 1 mm. Lower panels represent zoom images of the hippocampus, scale bar 200 µm. (**b,c**) Lysates from primary cultures of hippocampal neurons prepared from either *Dnm1*^+/+^ and *Dnm1*^+/R237W^ embryos were prepared and blotted for common presynaptic proteins and dynamin-1 interaction partners. (**b**) Representative blots displays levels of Synaptotagmin-1 (Syt1), Amphiphysin-1 (Amph1), C-src, Endophilin, Syndapin and Eps15. (**c**) Quantification of protein levels normalised to *Dnm1*^+/+^ ± SEM (n=3 independent cultures for all, all ns, Mann-Whitney test). (**d,e**) Workflow of quantitative proteomic analysis. Total protein content was cleaned onto SDS-PAGE gel before tryptic digestion. Proteins were analysed by high-resolution tandem MS. (**e**) Volcano plot displays 4237 quantified proteins, 39 which were depleted in Dnm1^+/R237W^ (blue), while 151 were enriched (red). Dnm1 level is displayed in green. (**f**) STRING network analysis of up and down regulated proteins in Dnm1^+/R237W^ synaptosomes. The resulting sub-complexes were subjected to MCL clustering at granulosity 4, which results in 18 clusters including 2 clusters with an average superior to 4 for the up-regulated proteins (proteasome core complex, ErbB signaling pathways) and 6 clusters including 2 clusters with an average superior to 4 for the down-regulated proteins (GPCR signaling pathway, dynactin complex).

Western blotting for common SV recycling proteins and dynamin-1 interaction partners in primary hippocampal neurons from *Dnm1*^+/R237W^ mice revealed no differences in their protein levels (**Figure 2b,c**). This was also the case in brain lysates from either 3 week or 6 week *Dnm1*^+/R237W^ mice (**Extended Data Figure 2c–g**). To determine more global changes in presynaptic protein expression, quantitative mass spectrometry was performed on nerve terminals isolated from either *Dnm1*^+/+^ or *Dnm1*^+/R237W^ littermates (**Figure 2d**). We established a list of 4237 quantified proteins associated to synapses, mitochondria and vesicular structures and transport (**Extended Data Figure 3**). This revealed 151 proteins that were significantly increased in *Dnm1*^+/R237W^ nerve terminals, with 39 significantly decreased (**Figure 2e, Extended Data Table 1**). To identify cellular functions that may be either upregulated or perturbed, network analysis using the STRING web tool (v.11.5) was performed on the proteins that were significantly altered. Upregulated proteins in *Dnm1*^+/R237W^ nerve terminals clustered around cell functions such as the proteasome core complex and the ErbB signalling pathway, whereas downregulated proteins were associated with G-protein coupled receptor signalling pathway and the dynactin complex (**Figure 2f**). Therefore, while there are no gross changes in architecture or protein expression in *Dnm1*^+/R237W^ mice, subtle variations are present at their nerve terminals that may reflect either disrupted cell signalling or potential compensatory mechanisms.

**Figure 3.**
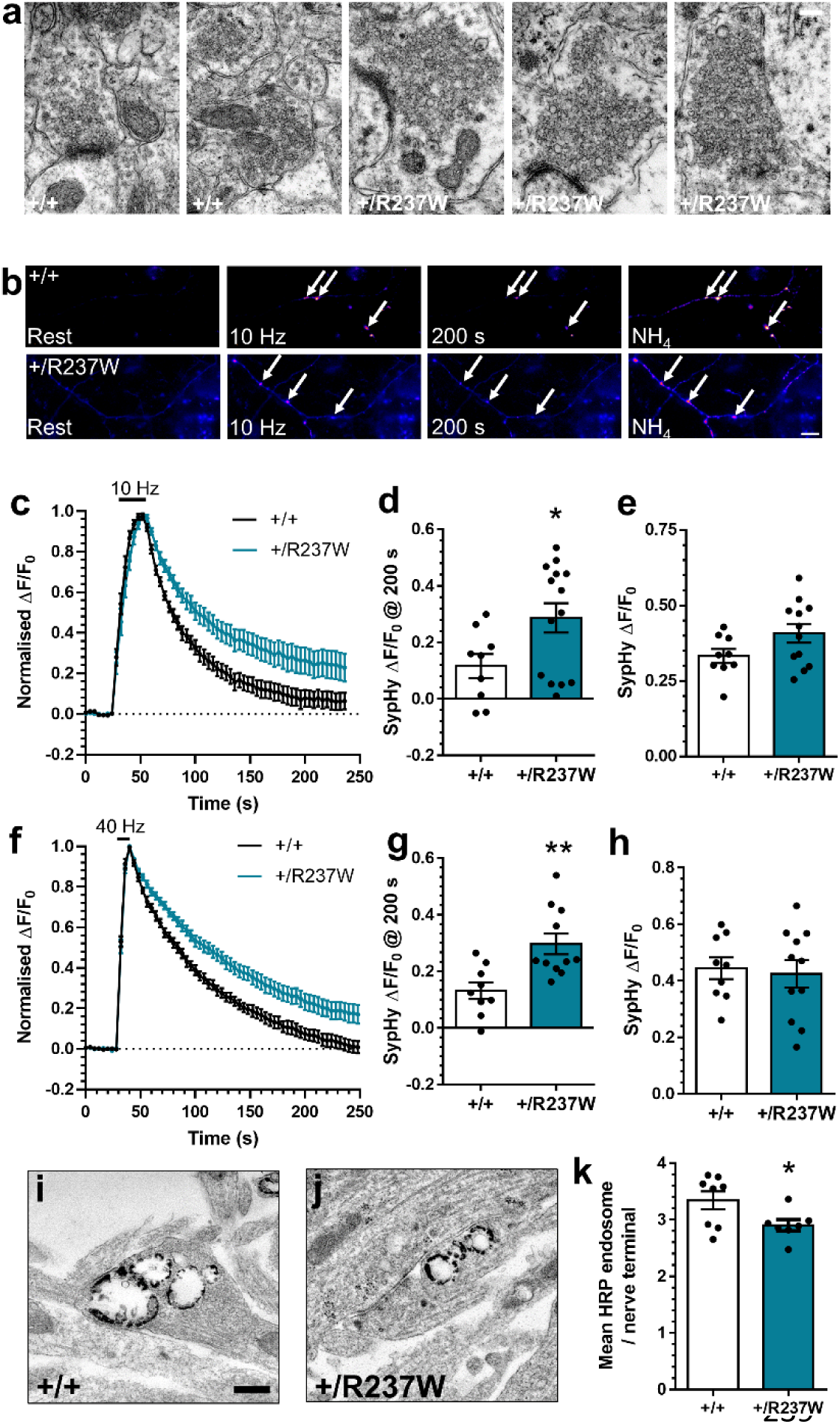
– *Dnm1*^+/R237W^ *neurons display dysfunctional SV endocytosis.* (**a**) Brains from 2 month-old *Dnm1*^+/+^ and *Dnm1*^+/R237W^ mice were perfusion fixed and processed for electron microscopy. Representative images reveal enlarged endosomes in *Dnm1*^+/R237W^ excitatory hippocampal nerve terminals, scale bar 250 nm. (**b-h**) Primary cultures of hippocampal neurons prepared from either *Dnm1*^+/+^ and *Dnm1*^+/R237W^ embryos were transfected with synaptophysin-pHluorin (sypHy) between 7-9 DIV. At 13-15 DIV, cultures were stimulated with a train of either (**c-e**) 300 action potentials (10 Hz) or (**f-h**) 400 action potentials (40 Hz). Cultures were pulsed with NH_4_Cl imaging buffer 180 s after stimulation. (**b**) Representative images of the sypHy response in *Dnm1*^+/+^ and *Dnm1*^+/R237W^ neurons are displayed at Rest, during 10 Hz stimulation, at 200 s and during NH_4_Cl. Arrows indicate responsive nerve terminals. Scale bar 10 μm. (**c,f**) Average sypHy response (ΔF/F_0_ ± SEM) normalised to the stimulation peak. (**d,g**) Average level of sypHy fluorescence (ΔF/F_0_ ± SEM) at 200 s (**d** Mann-Whitney test, n=9 *Dnm1*^+/+^, n=12 *Dnm1*^+/R237W^ *p=0.045; **g** Unpaired t test, n=9 *Dnm1*^+/+^, n=12 *Dnm1*^+/R237W^ **p=0.003). (**e,h**) Peak level of sypHy fluorescence (ΔF/F_0_ ± SEM) normalised to the NH_4_Cl challenge (**e** Unpaired t test, n=9 *Dnm1*^+/+^, n=12 *Dnm1*^+/R237W^ p=0.237; **h** Unpaired t test, n=9 *Dnm1*^+/+^, n=12 *Dnm1*^+/R237W^ p=0.751). (**i-k**) *Dnm1*^+/+^ and *Dnm1*^+/R237W^ neurons were stimulated with a train of 400 action potentials (40 Hz) in the presence of 10 mg/ml HRP. Representative images display HRP-labelled endosomes in *Dnm1*^+/+^ (**i**) and *Dnm1*^+/R237W^ (**j**) nerve terminals, scale bar 250 nm. (**k**) Average number of HRP-labelled endosomes per nerve terminal ± SEM (Unpaired t test, n=8 *Dnm1*^+/+^, n=7 *Dnm1*^+/R237W^ * p=0.039).

When hippocampal brain sections of *Dnm1*^+/R237W^ mice were examined at the ultrastructural level, their nerve terminals repeatedly displayed misshapen SVs and endosomal-like compartments, in contrast to *Dnm1*^+/+^ controls (**Figure 3a**). To determine whether this morphological phenotype resulted from a deficit in SV endocytosis, primary hippocampal cultures from *Dnm1*^+/R237W^ mice and *Dnm1*^+/+^ littermates were prepared. As before, SV exocytosis and endocytosis were monitored using the sypHy reporter (**Figure 3b**). Neurons from *Dnm1*^+/R237W^ mice displayed a significant slowing in SV endocytosis during challenge with two different stimulus trains (300 action potentials at 10 Hz or 400 action potentials at 40 Hz, **Figure 3c,d,f,g**). Furthermore, there was no significant effect on SV exocytosis during either stimulus train (**Figure 3e,h**). This was also the case when SV exocytosis was isolated in the presence of bafilomycin A1, which prevents acidification of SVs after endocytosis, removing contamination from retrieving SVs ^15^ (**Extended Data Figure 4a-d**). Therefore *Dnm1*^+/R237W^ neurons display a specific defect in SV endocytosis across a range of stimulus frequencies.

**Figure 4.**
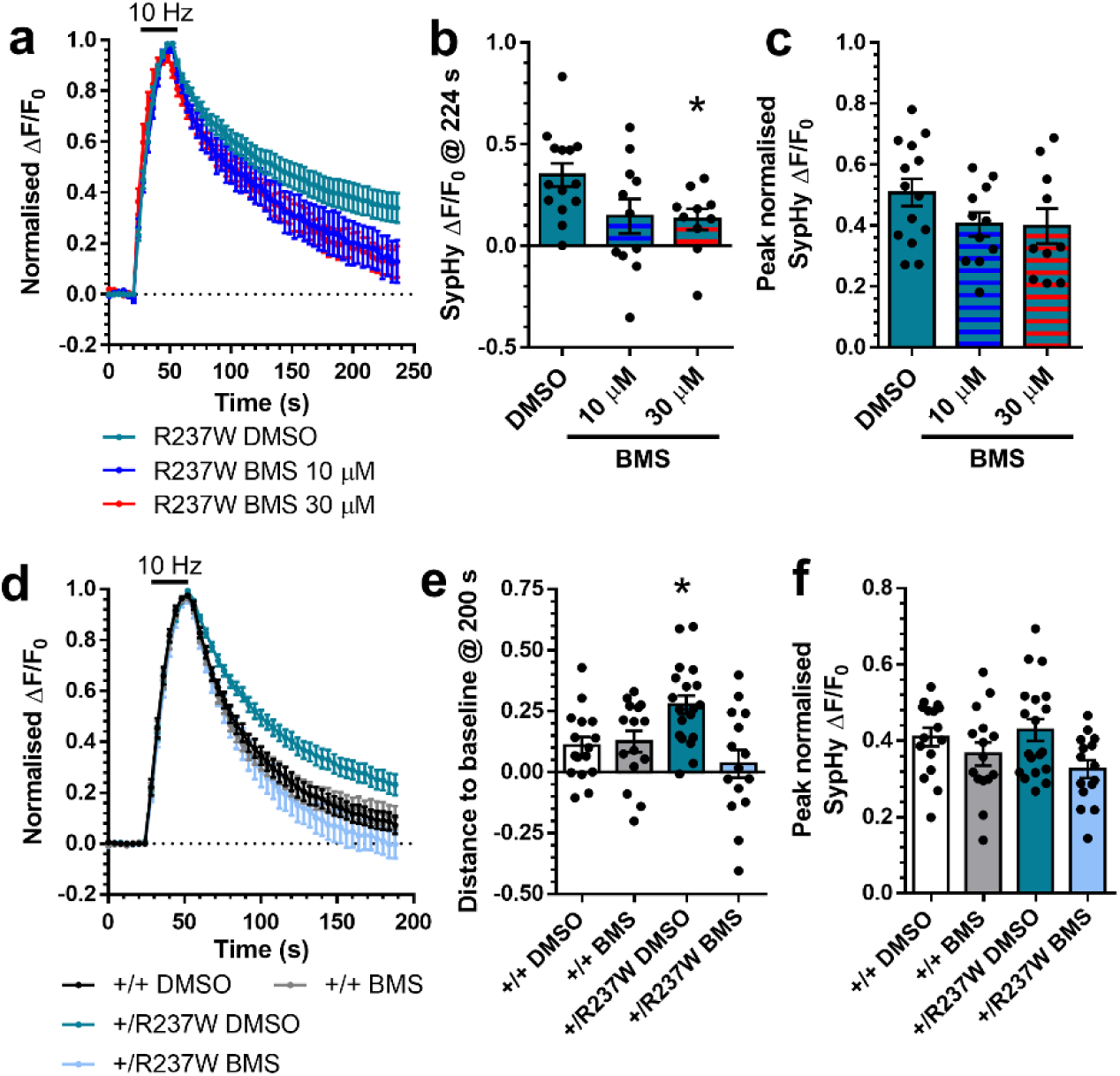
– *Dnm1*^+/R237W^ *mice display dysfunctional excitatory neurotransmission.* Neurotransmission at CA3/CA1 synapses was monitored using whole-cell patch clamp recording in acute hippocampal slices from *Dnm1*^+/+^ and *Dnm1*^+/R237W^ mice. (**a**) Example mEPSC events. Average frequency (**b**) and amplitude (**c**) of mEPSC events ± SEM (Mann-Whitney test, n=10 *Dnm1*^+/+^, n=12 *Dnm1*^+/R237W^, **b** p=0.0008, **c** p=0.665). (**d**) Paired pulse ratio (PPR) of evoked EPSCs as a function of the inter-stimulus interval (10-300 ms) ± SEM (Two-way ANOVA with Fishers LSD, n=11 *Dnm1*^+/+^, n=10 *Dnm1*^+/R237W^, ****p<0.0001). (**e**) Acute hippocampal slices were stimulated at a range of intensities (25, 50, 75, and 100 µA, 3 repeats at each intensity, frequency 0.05 Hz) in a pseudo random order. Evoked EPSC amplitude ± SEM is displayed (Two-way ANOVA with Fishers LSD, n=34 *Dnm1*^+/+^, n=39 *Dnm1*^+/R237W^, ****p<0.018, *p=0.043 50 µA, **p=0.005 75 µA, ***p=0.0004 100 µA). (**f,g**) Slices were stimulated with 600 APs (40 Hz). Average evoked EPSC amplitude normalized to peak response is displayed ± SEM (Two-way ANOVA, n=12 *Dnm1*^+/+^, n=13 *Dnm1*^+/R237W^, **p<0.0043). (**g**) Linear regression on the last 1 s of the cumulative EPSC plot in f (Two-way ANOVA, ****p<0.0001). (**h,j**) The average rate of readily releasable pool (RRP) replenishment (**h**) and mean RRP size (**j**) ± SEM were estimated from the linear regression plot in **g** (**h** Mann-Whitney test, **p=0.0045; **j** Unpaired t test, *p=0.0229). (**i**) Pr was calculated by dividing the amplitude of the first evoked EPSC by the effective RRP size (Unpaired t test, p=0.739).

To confirm the endocytosis defect via a complementary approach, morphological analysis was performed using activity-dependent uptake of the fluid phase marker horse radish peroxidase (HRP). After its accumulation, HRP can be converted to an electron dense product, to reveal the number of endocytic intermediates generated during stimulation ^16^. Recent studies have revealed that the majority of intermediates formed directly from the presynaptic plasma membrane are endosomes, which then shed SVs to refill the recycling pool ^17, 18^. *Dnm1*^+/R237W^ neurons displayed a significant reduction in HRP-labelled endosomes compared to *Dnm1*^+/+^ controls (**Figure 3i–k**), with no change in HRP endosome size (**Extended Data Figure 4e**). Therefore *Dnm1*^+/R237W^ neurons have an intrinsic and specific deficit in SV endocytosis.

### *Dnm1*^+/R237W^ mice display altered excitatory neurotransmission

SV endocytosis is essential to sustain neurotransmitter release ^19, 20^, therefore we predicted that the endocytosis defects observed in *Dnm1*^+/R237W^ neurons would translate into dysfunctional neurotransmission. To determine this, we examined neurotransmission at the excitatory Schaffer Collateral CA3-CA1 synapse, using whole-cell patch clamp recordings. We first determined the intrinsic properties of *Dnm1*^+/R237W^ neurons (**Extended Data Table 2**). Most parameters were unaffected, however differences in both Tau (membrane decay time) and capacitance (cell size) were detected. Alterations in capacitance may reflect dysfunctional endocytosis, whereas the elevated Tau value suggests the plasma membrane is slower to charge, reflecting a decrease in cell excitability. This increase in Tau is consistent with the increased half-width of action potentials in *Dnm1*^+/R237W^ neurons, in addition to the decay rate and rise time (**Extended Data Table 2**). Therefore, *Dnm1*^+/R237W^ neurons display alterations in their intrinsic excitability, which may be an adaptation to defects at the cell or circuit level.

The frequency, but not amplitude, of miniature excitatory postsynaptic currents (mEPSCs) was significantly reduced at *Dnm1*^+/R237W^ synapses (**Figure 4a–c**), suggesting a presynaptic SV recycling defect with no obvious postsynaptic phenotype. To confirm this, we determined whether evoked neurotransmission was impacted in *Dnm1*^+/R237W^ mice. Stimulation of CA3 axons with pairs of pulses at a range of inter-stimulus intervals revealed a significant decrease in the paired-pulse EPSC ratio in *Dnm1*^+/R237W^ synapses when compared to *Dnm1*^+/+^ (**Figure 4d**), suggesting an increased release probability (Pr) at these synapses. Furthermore, *Dnm1*^+/R237W^ synapses displayed an increase in evoked EPSC amplitudes across a range of stimulus intensities (**Figure 4e**). Thus, *Dnm1*^+/R237W^ synapses appear to have increased excitatory neurotransmission across a range of stimuli.

We next investigated whether *Dnm1*^+/R237W^ circuits could sustain neurotransmission during a prolonged train of action potential stimulation delivered at high frequency (600 APs at 40 Hz). *Dnm1*^+/R237W^ synapses displayed an inability to support neurotransmission during the stimulus train, when compared to *Dnm1*^+/+^ controls (**Figure 4f**). This finding, when considered with the reduced paired-pulse ratio, could be due to either increased Pr, or an inability to replenish SV pools. To determine this, the amplitude of the first evoked EPSC was divided by the effective readily releasable pool (RRP) size, estimated from the 40 Hz action potential train ^21^ (**Figure 4g**). Pr was unaffected in *Dnm1*^+/R237W^ circuits, whereas both the size and replenishment rate of the RRP was significantly reduced (**Figure 4h–j**). Therefore, excitatory neurotransmission in *Dnm1*^+/R237W^ circuits is initially augmented (most likely via increased Pr), however this enhancement cannot be sustained during an action potential train (most likely via reduced SV endocytosis), resulting in its depression.

### *Dnm1*^+/R237W^ mice display altered local field potential (LFP) activity and myoclonic jumping

Heterozygous mutations in the *DNM1* gene cause epileptic encephalopathies ^1, 2^. However many preclinical rodent models of monogenic epilepsies and neurodevelopmental disorders do not recapitulate the seizure activity observed in individuals with these disorders ^22^. When examined, *Dnm1*^+/R237W^ mice do not display overt spontaneous tonic-clonic seizures, however they do display “myoclonic jumping”, which involves bursts of highly active jumping (**Figure 5a, Extended Data Movie 1**). Importantly, *in vivo* electrophysiological recordings from *Dnm1*^+/R237W^ mice during these myoclonic jumps revealed increased generalised spiking activity during these events (**Figure 5b**). This appears not to be a consequence of jumping *per se*, since this activity was far more pronounced in *Dnm1*^+/R237W^ mice when compared to the rare jumping events in *Dnm1*^+/+^ mice (**Figure 5b**). This suggests that this behaviour is a consequence of *bone fide* seizure activity. Therefore, the *Dnm1*^+/R237W^ mouse has both construct and face validity as a preclinical model of *DNM1* epileptic encephalopathy.

**Figure 5.**
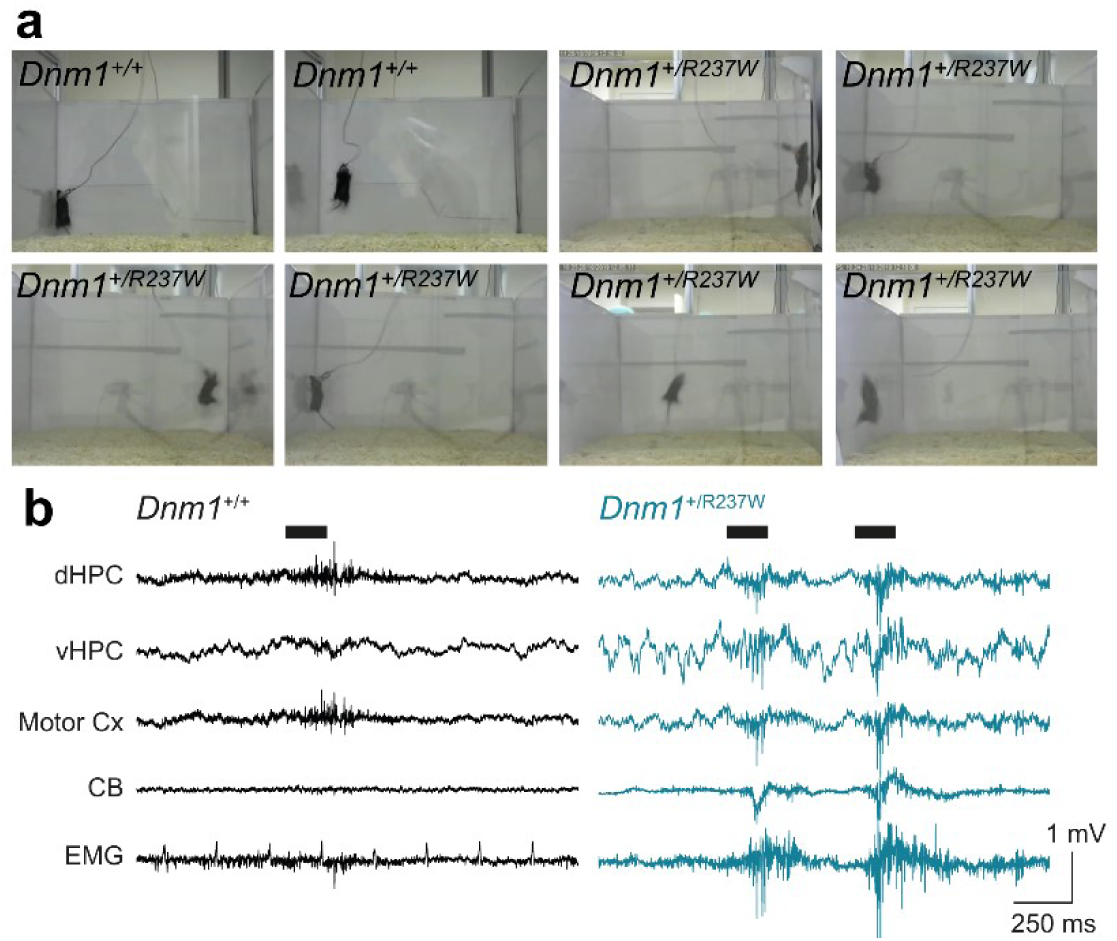
– Dnm1^+/R237W^ mice display myoclonic jumping seizure-like activity. (**a**) Still images displaying the typical jumping behaviour of both *Dnm1*^+/+^ and *Dnm1*^+/R237W^ mice. *Dnm1*^+/+^ mice occasionally jumped, however *Dnm1*^+/R237W^ mice displayed stereotypical and burst-like events. (**b**) Example traces of *in vivo* LFP and electromyogram (EMG) recordings from dorsal hippocampus (dHPC, ventral hippocampus (vHPC), motor cortex (Motor Cx) and cerebellum (CB) during jumping activity in *Dnm1*^+/+^ and *Dnm1*^+/R237W^ mice.

### Reversal of presynaptic and circuit phenotypes in *Dnm1*^+/R237W^ neurons by BMS-204352

The presynaptic and circuit phenotypes observed in *Dnm1*^+/R237W^ neurons are strongly supportive of dysfunctional SV endocytosis being the key driver of the myoclonic jumping observed in *Dnm1*^+/R237W^ mice. Therefore, we next determined whether correction of SV endocytosis could restore normal neurotransmission and ablate the observed behavioural phenotypes. The small molecule BMS-20352 ((3S)-(+)-(5-chloro-2-methoxyphenyl)-1,3-dihydro-3-fluoro-6-(trifluoromethyl)-2H-indol-2-one) was chosen for this task since it is a therapeutic safe for use in humans ^23^ and can correct behavioural defects in a preclinical model of fragile X syndrome ^24^. This latter effect prompted us to investigate its action, since a number of fragile X syndrome model systems display circuit hyperexcitability ^25^. We first examined the effect of BMS-204352 in primary cultures of *Dnm1*^+/+^ hippocampal neurons overexpressing both Dyn1_WT_-mCer and sypHy. Intriguingly, a dose-dependent acceleration of SV endocytosis was observed at time points after stimulation, with a reduction in SV exocytosis also observed at the highest dose (**Extended Data Figure 5**). Therefore BMS-204352 may have the potential to correct presynaptic defects in *Dnm1*^+/R237W^ neurons.

BMS-204352 displays positive modulatory effects on both neuronal K_v_7 channels and BK channels, whereas it is a negative modulator of both K_v_7.1 channels and GABA_A_ receptors ^26, 27^. This spectrum of activity across multiple potassium channel subtypes suggest that it may accelerate SV endocytosis via a series of different mechanisms. To determine this, we examined SV endocytosis and exocytosis in *Dnm1*^+/+^ hippocampal neurons expressing sypHy in the presence of a series of potassium channel modulators. These were: two structurally-unrelated BK channel agonists (NS11021 and BMS-191011), a BK channel antagonist (Paxilline), a K_v_7 channel activator (Retigabine), and a K_v_7 channel inhibitor (XE-991). No modulator was able to accelerate SV endocytosis in the manner observed with BMS-204352 (**Extended Data Figure 6**). Therefore, the action of BMS-204352 on SV endocytosis is not due to modulation of a specific class of ion channels, suggesting its presynaptic effects are an amalgamation of the modulation of some or all of these channels, or an as yet unidentified off-target effect.

**Figure 6.**
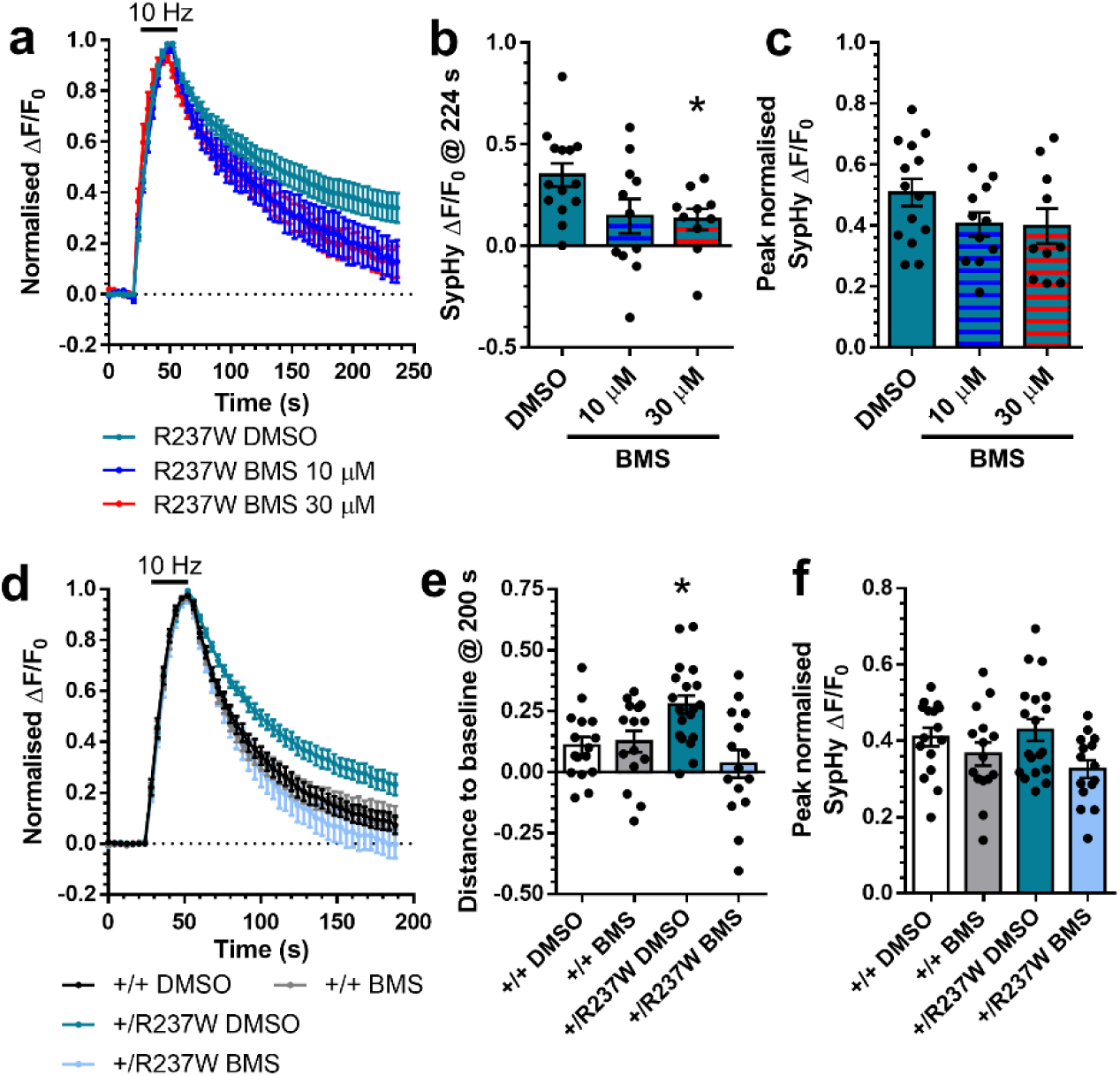
– *BMS-204352 corrects dominant negative effect of R237W mutation on SV endocytosis.* (**a-c**) Primary cultures of hippocampal neurons prepared from *Dnm1*^+/+^ embryos were transfected with synaptophysin-pHluorin (sypHy) and Dyn1_R237W_-mCer between 11-13 DIV. At 13-15 DIV, cultures were stimulated with a train of 300 action potentials (10 Hz) in the presence of either 10 μM or 30 μM BMS-204352 or a vehicle control (DMSO). Cultures were pulsed with NH_4_Cl imaging buffer 180 s after stimulation. (**a**) Average sypHy response (ΔF/F_0_ ± SEM) normalised to the stimulation peak. (**b**) Average level of sypHy fluorescence (ΔF/F_0_ ± SEM) at 224 s (One-way ANOVA, n=14 DMSO, N=11 10 μM, n=10 30 μM, *p=0.0467 DMSO vs 30 μM). (**c**) Peak level of sypHy fluorescence (ΔF/F_0_ ± SEM) normalised to the NH_4_Cl challenge (One-way ANOVA, n=10 DMSO, N=8 10 μM, n=6 30 μM, all ns). (**e-f**) Primary cultures of hippocampal neurons prepared from either *Dnm1*^+/+^ or *Dnm1*^+/R237W^ embryos were transfected with sypHy between 7-9 DIV. At 13-15 DIV, cultures were stimulated with a train of 300 action potentials (10 Hz) in the presence of either 30 μM BMS-204352 or a vehicle control (DMSO). Cultures were pulsed with NH_4_Cl imaging buffer 180 s after stimulation. (**d**) Average sypHy response (ΔF/F_0_ ± SEM) normalised to the stimulation peak. (**e**) Average level of sypHy fluorescence (ΔF/F_0_ ± SEM) at 200 s (One-way ANOVA, n=19 DMSO *Dnm1*^+/+^, n=15 BMS *Dnm1*^+/+^, n=16 DMSO *Dnm1*^+/R237W^, n=15 BMS *Dnm1*^+/R237W^, *p=0.0183 DMSO *Dnm1*^+/+^ vs DMSO *Dnm1*^+/R237W^). (**f**) Peak level of sypHy fluorescence (ΔF/F_0_ ± SEM) normalised to the NH_4_Cl challenge (One-way ANOVA n=19 DMSO *Dnm1*^+/+^, n=15 BMS *Dnm1*^+/+^, n=16 DMSO *Dnm1*^+/R237W^, n=15 BMS *Dnm1*^+/R237W^, all ns).

Regardless of the BMS-204352 mechanism of action, we next examined whether it was able to correct defective SV endocytosis due to expression of the R237W dynamin-1 mutant. We first determined its effect on *Dnm1*^+/+^ cells overexpressing Dyn1_R237W_-mCer. In these neurons, BMS-204352 fully restored SV endocytosis kinetics (**Figure 6a,b**), suggesting it may be a viable intervention to correct dysfunction in *Dnm1*^+/R237W^ neurons. When the effect of BMS-204352 on SV endocytosis was examined in primary cultures of *Dnm1*^+/R237W^ neurons, a full correction of SV endocytosis kinetics was again observed when compared to *Dnm1*^+/+^ neurons (**Figure 6d,e**). BMS-204352 had no significant effect on SV exocytosis in either *Dnm1*^+/+^ neurons with overexpressed Dyn1_R237W_-mCer, or *Dnm1*^+/R237W^ neurons (**Figure 6c,f**). Therefore BMS-204352 restores SV endocytosis that was previously rendered dysfunctional via the mutant R237W *Dnm1* allele.

We next determined whether BMS-204352 could correct the observed dysfunction of excitatory neurotransmission in *Dnm1*^+/R237W^ mice, since we predicted that defects in circuit activity were a result of impaired SV endocytosis. When applied to *Dnm1*^+/+^ hippocampal slices, BMS-204352 had no effect on evoked EPSC amplitudes across a range of stimuli (**Figure 7a**). However, BMS-204352 fully restored normal evoked EPSC amplitudes in *Dnm1*^+/R237W^ slices to *Dnm1*^+/+^ levels, across the same range of stimulus intensities (**Figure 7b**). Therefore BMS-204352 can restore normal evoked excitatory neurotransmission in *Dnm1*^+/R237W^ circuits.

**Figure 7.**
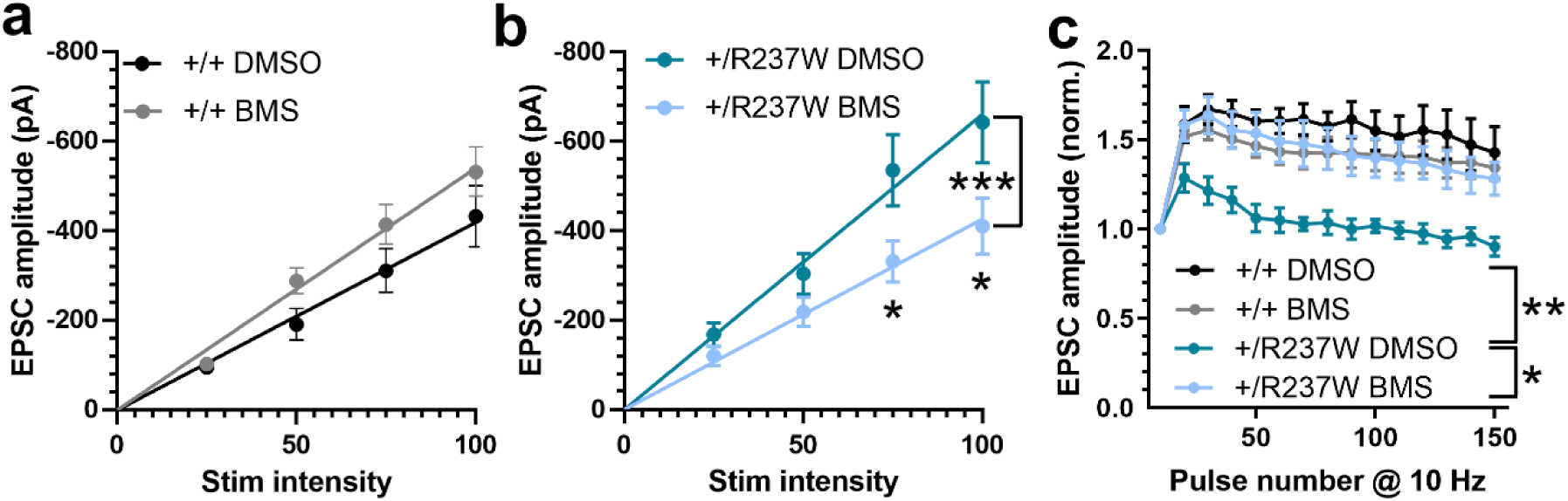
– *BMS-204352 corrects neurotransmission defects in Dnm1*^+/R237W^ *mice.* (**a,b**) Neurotransmission at CA3/CA1 synapses was monitored in acute hippocampal slices from either *Dnm1*^+/+^ (**a**) or *Dnm1*^+/R237W^ (**b**) mice. Slices were stimulated at a range of intensities (25, 50, 75, and 100 µA, 3 repeats at each intensity, frequency 0.05 Hz) in a pseudo random order in the presence of 30 μM BMS-204352 or a vehicle control (DMSO). Evoked EPSC amplitude ± SEM is displayed (Two-way ANOVA with Sidak’s multiple comparison test, **a** n=13 DMSO, n=11 BMS, all ns; **b** n=13 DMSO, n=13 BMS, ***p=0.0004, *p=0.041 75 μA, *p=0.0145 100 μA). (**c**) Evoked EPSC amplitude for *Dnm1*^+/+^ or *Dnm1*^+/R237W^ hippocampal slices stimulated with a 10 AP train (10 Hz, normalized to first pulse) in the presence of either 30 μM BMS-204352 or a vehicle control (DMSO). Two-way ANOVA with Dunnett’s multiple comparison test, n=5 DMSO *Dnm1*^+/+^, n=6 BMS *Dnm1*^+/+^, n=5 DMSO *Dnm1*^+/R237W^, n=7 BMS *Dnm1*^+/R237W^, **p=0.002 DMSO *Dnm1*^+/+^ vs DMSO *Dnm1*^+/R237W^, *p=0.013 DMSO *Dnm1*^+/R237W^ vs BMS *Dnm1*^+/R237W^.

We next examined whether BMS-204352 could reverse short-term plastic changes in excitatory neurotransmission by monitoring synaptic facilitation evoked via a 10 Hz AP train (15 sec). In *Dnm1*^+/+^ slices, a pronounced facilitation was observed (**Figure 7c**), in agreement with previous studies ^28^. In contrast, no facilitation of excitatory neurotransmission was observed in *Dnm1*^+/R237W^ slices (**Figure 7c**). Application of BMS-204352 to *Dnm1*^+/R237W^ hippocampal slices fully restored facilitation to wild-type levels and had no effect on the *Dnm1*^+/+^ response (**Figure 7c**). Therefore BMS-204352 corrects fundamental defects in evoked excitatory neurotransmission and short-term plasticity in *Dnm1*^+/R237W^ circuits.

Finally, we determined whether BMS-204352 was able to correct the myoclonic jumping phenotype in *Dnm1*^+/R237W^ mice. *Dnm1*^+/R237W^ mice and *Dnm1*^+/+^ littermate controls were habituated in an open field arena for 30 minutes on day 1. This protocol was repeated for 5 days. On days 2 and 4, mice were dosed with either BMS-204352 or a vehicle control in a counterbalanced manner (**Figure 8a**). Mice were also monitored on days 3 and 5 to examine baseline behaviour (Washout, **Figure 8a**). When baseline behaviour of *Dnm1*^+/+^ and *Dnm1*^+/R237W^ mice were examined, robust differences in both the number of myoclonic jumps and bursts of jumps (defined as a train of at least 2 myoclonic jumps with less than 2s between consecutive jumps) were observed (**Extended Data Figure 7a,b**). This phenotype was not due to increased general activity, since there is no significant change in the distance travelled by *Dnm1*^+/R237W^ when compared to *Dnm1*^+/+^ controls (**Extended Data Figure 7c**).

**Figure 8.**
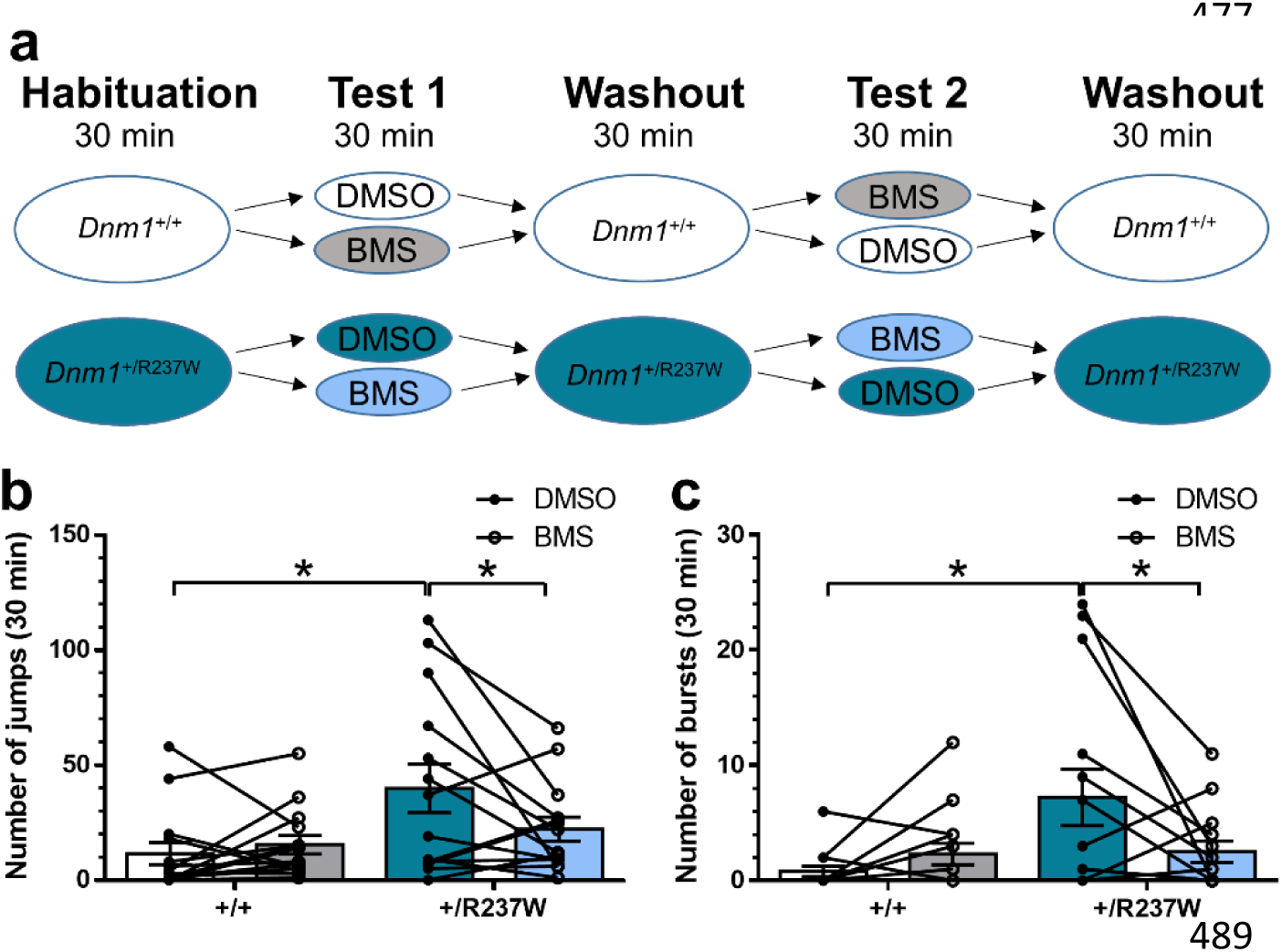
– *BMS-204352 corrects seizure-like events in Dnm1*^+/R237W^ *mice. Dnm1*^+/+^ and *Dnm1*^+/R237W^ mice were placed in an open field chamber for a 30 min period for 5 days. After habituation on day 1, mice were dosed with either BMS-204352 or a vehicle control (DMSO) on days 2 and 4, with drug washout on days 3 and 5. Delivery of drug treatment was interleaved between days 2 and 4. (**a**) Schematic of the experimental protocol. (**b,c**) Average number of myoclonic jumps (**b**) or bursts (**c**). (**b**) General linear model (repeated measures) with Bonferroni multiple comparisons n=14 for all, * p=0.019 DMSO *Dnm1*^+/R237W^ vs BMS *Dnm1*^+/R237W^, * p=0.021 DMSO *Dnm1*^+/+^ vs DMSO *Dnm1*^+/R237W^, all other ns (**c**) General linear model (repeated measures) n=14 for all, * p=0.016 DMSO *Dnm1*^+/+^ vs DMSO *Dnm1*^+/R237W^ * p=0.011 DMSO *Dnm1*^+/R237W^ vs BMS *Dnm1*^+/R237W^ all other ns.

During the test phase, the phenotypes that were observed during washout were retained in vehicle-treated *Dnm1*^+/+^ and *Dnm1*^+/R237W^ mice. Specifically, vehicle-treated *Dnm1*^+/R237W^ mice displayed a significant increase in the total number of myoclonic jumps (**Figure 8b**) and the number of jumping bursts (**Figure 8c**) when compared to *Dnm1*^+/+^ controls. Delivery of BMS-204352 to *Dnm1*^+/+^ mice had no significant effect on these parameters (**Figure 8b,c**). In contrast, BMS-204352 fully corrected both jumping phenotypes in *Dnm1*^+/R237W^ mice to the levels observed in *Dnm1*^+/+^ mice (**Figure 8b,c**). Importantly, this correction was not due to depression of locomotive activity, since the distance travelled was not significantly different when *Dnm1*^+/R237W^ mice treated with or without BMS-204352 were compared (**Extended Data Figure 7c**). Furthermore, BMS-204352 had no effect on the time spent in the middle of the open area, indicating that the correction of seizure phenotypes was not due to previously documented anxiolytic effects of the molecule ^27^ (**Extended Data Figure 7d**). In summary, these results suggest that BMS-204352 has high potential for therapy in *DNM1* epileptic encephalopathy, since it corrects dysfunction at the cellular, circuit and behavioural level in a novel preclinical model of this disorder.

## Discussion

Heterozygous *DNM1* mutations are responsible for a novel form of epileptic encephalopathy ^1, 2^. Here, we confirmed that the most common pathogenic *DNM1* mutation, R237W, disrupts dynamin-1 enzyme activity and SV endocytosis. Furthermore, using the *Dnm1*^+/R237W^ mouse, we revealed that dysfunctional SV endocytosis translates into altered excitatory neurotransmission and ultimately seizure-like phenotypes. Importantly, these phenotypes were corrected at the cell, circuit and *in vivo* level via the acceleration of SV endocytosis using BMS-204352. This study therefore provides the first direct link between dysfunctional SV endocytosis and epilepsy, but moreover reveals that SV endocytosis is a viable therapeutic route for monogenic intractable epilepsies.

### A novel model of *DNM1* epileptic encephalopathy

The R237W mutation was chosen for our preclinical model, since it is the most prevalent missense mutation in the *DNM1* gene (8 from 33 cases ^1, 2^). The *Dnm1*^+/R237W^ mouse appears to have both face and construct validity and therefore is predicted to be of high value for future therapeutic studies. These mice displayed a selective defect in SV endocytosis, excitatory neurotransmission and a characteristic jumping phenotype. This behavioural phenotype occurred co-incident with increased generalised spiking activity, providing evidence that it is precipitated via seizure-like events. We named this jumping phenotype “myoclonic jumping” since it appears similar to phenotypes in several preclinical epilepsy models observed either in isolation or in progression towards full tonic-clonic seizures ^29–31^. Furthermore, it is also observed in autism / neurodegeneration models as a measure of repetitive and stereotypic behaviour ^32, 33^.

One intriguing finding was the alteration in excitatory neurotransmission in *Dnm1*^+/R237W^ mice. Previous studies in *Dnm1*^-/-^ neurons had revealed a disproportionate effect on inhibitory neurotransmission, with large effects on evoked IPSCs when compared to EPSCs, and faster and more extensive depression of inhibitory neurotransmission during action potential trains ^34^. Furthermore, in *Dnm1/3*^-/-^ synapses, a strong facilitation of excitatory neurotransmission was observed during both low and high frequency stimulation, with mEPSC frequency, evoked EPSC amplitude, RRP size and Pr all decreased ^35^. *Dnm1*^+/R237W^ synapses also display reduced mEPSC frequency and decreased RRP, however in contrast, we observe an increase in both Pr and evoked EPSCs in addition to an absence of STP during action potential trains. Therefore, even when SV endocytosis is disrupted to a similar extent between *Dnm1*^+/R237W^ and *Dnm1*^-/-^ neurons, the dominant-negative R237W mutation exerts discrete effects on circuit activity not observed in models of loss of dynamin-1 function. Determining how *Dnm1*^+/R237W^ neurons modify brain circuit properties will be a key avenue for future study.

### Correction of phenotypes in *Dnm1*^+/R237W^ mice using BMS-204352

We observed a full correction of cellular, circuit and *in vivo* phenotypes via the delivery of BMS-204352. BMS-204352 was developed as a BK channel agonist for the treatment of stroke ^36^, however in Phase III trials it failed to display efficacy superior to placebo ^23^. Nevertheless, the drug exhibited an excellent safety profile, identifying it as a promising candidate for repurposing studies. Because BMS-204352 displayed both positive and negative modulatory effects against a number of potassium channel subtypes ^26, 27^, we employed a series of channel openers and blockers to elucidate its mechanism of action. Intriguingly, no drug from this palette of modulators recapitulated its observed modifying activity on SV endocytosis. Therefore BMS-204352 may have additional off-target effects responsible for its reversal of phenotypes in the *Dnm1*^+/R237W^ mouse. This question is under active investigation.

BMS-204352 is therefore a promising lead compound for future trials in *DNM1* epileptic encephalopathy. There is also potential for its use to be wider than this specific condition, since a cohort of monogenic neurodevelopmental disorders are predicted to have SV endocytosis defects at their core. For example, a series of frameshift, nonsense and missense mutations in essential SV endocytosis and SV cargo clustering genes have been identified in individuals with intellectual disability, autism and epilepsy ^22^, including the coat protein clathrin ^37^, adaptor protein complexes ^38^, SV cargo retrieval ^39–41^ and regulators of endocytosis such as TBC1D24 ^42, 43^. Furthermore, neurons derived from preclinical models for prevalent monogenic conditions such as fragile X syndrome and CDKL5 deficiency disorder have recently been discovered to display defects in SV retrieval ^16, 44^. With dysfunctional SV endocytosis emerging as a key convergence point in these monogenic conditions, upregulation of SV endocytosis via BMS-204352 may therefore provide a potential treatment to restore essential recycling mechanisms and normal function.

In conclusion, we have key cell, circuit and behavioural defects in a mouse model of *DNM1* epileptic encephalopathy, which provide important information on the molecular locus of seizure activity. Furthermore, an agent that accelerates SV endocytosis corrects all of these defects, suggesting intervention via this trafficking pathway is an exciting therapeutic route.

## Acknowledgements

This research was funded by grant awards to MAC (Epilepsy Research UK (P2003), Wellcome Trust Investigator Award (204954/Z/16/Z) and RS McDonald Fund) to MT (Wellcome Trust Multi-User Equipment grant (212947/Z/18/Z)) and AGS (Epilepsy Research UK (F1603)). For the purpose of open access, the author has applied a CC-BY public copyright license to any author accepted manuscript version arising from this submission. We thank Steven Mitchell for EM sample processing.

## Author Contributions

Conceptualization, KB, MAC; Methodology, KB, KLD, MP, MS, EB, AG, ECD, MT, AGS; Formal Analysis, KB, KLD, AG, ECD, MP, MS, AGS; Investigation, KB, KLD, AGS, MAC; Resources, MAC, MT, AGS; Writing – Original Draft, KB, MAC; Writing – Review & Editing, all authors; Funding Acquisition, MAC, MT, AGS.

## Methods

### Materials

Unless otherwise specified, all cell culture reagents were obtained from Invitrogen (Paisley, UK). Foetal bovine serum was from Biosera (Nuaille, France). Papain was obtained from Worthington Biochemical (Lakewood, NJ, USA). Caesium-gluconate, tetrodotoxin (TTX) and picrotoxin were from Hello Bio (Bristol, UK). Na_2_GTP was from Scientific Laboratory Supplied (Newhouse, UK), whereas Na_2_-creatine was from (Merck, London, UK). BMS-204352 was from Bio-Techne Ltd (Abingdon, UK). All other reagents were obtained from Sigma-Aldrich (Poole, UK) unless specified. Synaptophysin-pHluorin (sypHy) was provided by Prof. L. Lagnado (University of Sussex, UK). Rat dynamin-1aa fused to mCerulean at its C-terminus ^45^ was subjected to site-directed mutagenesis to generate both R237W (forward primer ATTGGCGTGGTGAAC<u>TGG</u>AGCCAGAAGGACATA, reverse primer TATGTCCTTCTGGCT<u>CCA</u>GTTCACCACGCCAAT) and K44A mutations (forward primer GGCCAGAGCGCCGGC<u>GCG</u>AGCTCGGTGCTGGAC, reverse primer CTCCAGCACCGAGCT<u>CGC</u>GCCGGCGCTCTGGCC). Base changes were confirmed by Source Bioscience Sanger Sequencing (Glasgow, UK).

### Generation of *Dnm1*^+/R237W^ mice

The *Dnm1*^+/R237W^ mouse was generated by Horizon Discovery (St. Louis, USA). Briefly, the codon encoding R237 within the *Dnm1* gene was targeted using CRISPR-Cas9 technologies on a C57Bl/6J genetic background. This resulted in the modification of the *Dnm1* gene sequence from CGGAGC (equivalent amino acids 237/238-RS) to TGGTCT (amino acids 237/238-WS). Mice were maintained as heterozygotes with the colony sustained via breeding with wild-type C57Bl/6J mice. Genotyping was performed by Transnetyx (Cordova, TN, USA).

Animal work was performed in accordance with the UK Animal (Scientific Procedures) Act 1986, under Project and Personal Licence authority and was approved by the Animal Welfare and Ethical Review Body at the University of Edinburgh (Home Office project licences – 7008878 and PP5745138 to Prof. Cousin and PP1538548 to Dr. Gonzalez-Sulser). Specifically, all animals were killed by Schedule 1 procedures in accordance with UK Home Office Guidelines; adults were killed by cervical dislocation or exposure to CO_2_ followed by decapitation, whereas embryos were killed by decapitation followed by destruction of the brain.

### Cell culture and transfections

Heterozygous *Dnm1*^+/R237W^ mice were mated with *Dnm1*^+/+^ mice to produce either *Dnm1*^+/+^ or *Dnm1*^+/R237W^ offspring. Hippocampi from each embryo were processed separately to avoid contamination across genotypes. Dissociated primary hippocampal cultures were prepared from embryos as previously described ^16^. Briefly, isolated hippocampi were digested in a 10 U/mL papain solution (Worthington Biochemical, LK003178) at 37°C for 20 min. The papain was then neutralised using DMEM F12 (ThermoFisher Scientific, 21331-020) supplemented with 10 % Foetal bovine serum (BioSera, S1810-500) and 1 % penicillin/streptomycin (ThermoFisher Scientific, 15140-122). Cells were triturated to form a single cell suspension and plated at 5 x 10^4^ cells per coverslip on laminin (10 µg/ mL; Sigma Aldrich, L2020) and poly-D-lysine (Sigma Aldrich, P7886) coated 25 mm glass coverslips (VWR International Ltd, Lutterworth, UK). Cultures were maintained in Neurobasal media (ThermoFisher Scientific, 21103-049) supplemented with 2 % B-27 (ThermoFisher Scientific, 17504-044), 0.5 mM L-glutamine (ThermoFisher Scientific, 25030-024) and 1% penicillin/streptomycin. After 2-3 days *in vitro* (DIV), 1 µM of cytosine arabinofuranoside (Sigma Aldrich, C1768) was added to each well to inhibit glial proliferation. Hippocampal neurons were transfected with synaptophysin-pHluorin (sypHy) and/or Dyn1-mCer using Lipofectamine 2000 (ThermoFisher Scientific, 11668027) as per manufacturer’s instructions and imaged at *DIV* 13-15.

### Imaging of SV recycling using sypHy

Imaging of SV recycling was monitored using sypHy as previously described ^16^. SypHy-transfected hippocampal cultures were mounted in a Warner Instruments (Hamden, CT, USA) imaging chamber with embedded parallel platinum wires (RC-21BRFS) and were mounted on a Zeiss Axio Observer D1 inverted epifluorescence microscope (Cambridge, UK). Neurons were challenged with field stimulation using a Digitimer LTD MultiStim system-D330 stimulator (current output 100 mA, current width 1 ms) either at 10 Hz for 30 s or 40 Hz for 10 s. Neurons were visualised at 500 nm band pass excitation with a 515 nm dichroic filter and a long-pass >520 nm emission filter, with images captured using an AxioCam 506 mono camera (Zeiss) with a Zeiss EC Plan Neofluar 40×/1.30 oil immersion objective. Image acquisition was controlled using Zen Pro software (Zeiss). Imaging time courses were acquired at 4 s intervals while undergoing constant perfusion with imaging buffer (119 mM NaCl, 2.5 mM KCl, 2 mM CaCl_2_, 2 mM MgCl_2_, 25 mM HEPES, 30 mM glucose at pH 7.4, supplemented with 10 μM 6-cyano-7-nitroquinoxaline-2,3-dione (Abcam, Cambridge, UK, ab120271) and 50 μM DL-2-Amino-5-phosphonopentanoic acid (Abcam, Cambridge, UK, ab120044). Alkaline buffer (50 mM NH_4_Cl substituted for 50 mM NaCl) was used to reveal the maximal pHluorin response. SV fusion during stimulation was measured by stimulating sypHy-transfected neurons (10 Hz, 90 s) in the presence of 1 µM bafilomycin A1 (Cayman Chemical Company, Ann Arbor Michigan, USA, 11038). BMS-204352 and other potassium channel modulators in imaging buffer were perfused over neurons 2 min prior to and during imaging until addition of alkaline buffer.

Time traces were analysed using the FIJI distribution of Image J (National Institutes of Health). Images were aligned using the Rigid body model of the StackReg plugin (https://imagej.net/StackReg). Nerve terminal fluorescence was measured using the Time Series Analyser plugin (https://imagej.nih.gov/ij/plugins/time-series.html). Regions of interest (ROIs) 5 pixels in diameter were placed over nerve terminals that responded to the electrical stimulus. A response trace was calculated for each cell by averaging the individual traces from each selected ROI. Inhibition of SV endocytosis was calculated as remaining fluorescence 140 s after termination of stimulation.

### HRP uptake in hippocampal cultures

Hippocampal cultures were mounted in the RC-21BRFS stimulation chamber and challenged with 400 action potentials (40 Hz) in the presence of 10 mg/ml HRP (Sigma Aldrich, P8250) supplemented imaging buffer. Immediately following the end of stimulation, cultures were washed in imaging buffer to remove non-internalised HRP and fixed with a solution of 2 % glutaraldehyde (Electron Microscopy Sciences, Hatfield, USA, 16019) and 2% PFA in 0.1 M in phosphate buffer (PB). After washing in 0.1 M PB, HRP was developed with 0.1 % 3,3’-diaminobenzidine (Fluka Chemica, Gillingham, UK, 22204001) and 0.2 % v/v hydrogen peroxide (Honeywell, Muskegon, USA, 216763) in PB. After further washing in PB, cultures were stained with 1 % osmium tetroxide (TAAB laboratory and microscopy, Aldermaston, UK, O015/1) for 30 min. Samples were then dehydrated using an ethanol series and polypropylene oxide (Electron Microscopy Sciences, Hatfield, USA, 20411) and embedded using Durcupan resin (Sigma Aldrich, 44610). Samples were sectioned, mounted on grids, and viewed using an FEI Tecnai 12 transmission electron microscope (Oregon, USA). Intracellular structures that were <61 nm in diameter were arbitrarily designated to be SVs, whereas larger structures were considered endosomes. The area of individual endosomes was obtained by tracing the circumference using the freehand selections tool in ImageJ and measuring the resulting area. Typically, 20 fields of view were acquired for one coverslip of cells. In nerve terminals that contained HRP, the average number of HRP-labelled endosomes and SVs per nerve terminal was calculated for each coverslip and represents the experimental n.

### Immunocytochemistry

Immunofluorescence staining and analysis was performed as previously described ^16^. Briefly, hippocampal neurons were fixed with 4% paraformaldehyde (PFA) in PBS for 15 min at room temperature. PFA was then removed and cells were quenched 2×5 min with 50 mM NH_4_Cl in PBS. Cells were then washed 4× 5 min with PBS. Before staining, cells were permeabilised in 1% bovine serum albumin (BSA) in PBS-Triton 1% for 5 min. Cells were then washed in PBS before blocking in 1% BSA in PBS at room temperature for 1 h. After blocking, cells were left to incubate in primary antibody diluted in blocking solution for 30-45 min (chicken anti-GFP (Abcam ab13970) 1:500; rabbit anti-SV2A (Abcam 32942) 1:200; goat anti-dynamin-1 (Santa Cruz sc-6402) 1:200). Following 4 x 5 min washes, cells were left to incubate in secondary antibody (goat anti-chicken Alexa-Fluor-488 (Invitrogen A11039) 1:1000; goat anti-rabbit Alexa-Fluor-568 (Invitrogen A21069) 1:1000; donkey anti-goat Alexa-Fluor-647 (Invitrogen A21447) 1:1000) diluted in blocking buffer for 30-45 min at room temperature in the dark. After washing, coverslips were mounted to slides using FluorSave Reagent. Alexa Fluor 488 and 568 images were acquired using a dual camera imaging system (Zeiss). The signal was filtered by a double band pass excitation filter (470/27 + 556/25) with beam splitter (490 + 575) and emission filters 512/30 and 630/98 (Zeiss) respectively. Alexa Fluor 647 was visualised with a 640 nm excitation and a 690/50 band pass emission filter. For each image analysed, ROIs were placed over the transfected neuron, a non-transfected neuron and the background. This allowed measurement of levels of overexpression of mCer-Dyn1 within neurons on the same coverslip by comparing the overexpression to normal expression levels. Background fluorescence was subtracted from all signals. For each coverslip, 4-6 fields with transfected neurons were acquired. The n is the number of transfected cells imaged.

### Immunohistochemistry

Two-month-old *Dnm1*^+/+^ or *Dnm1*^+/R237W^ male mice were administered a lethal dose of sodium pentobarbital and transcardially perfused with cold PBS followed by cold PFA (PFA, 4% in 0.1M PB). Brains were extracted and fixed for 24 hours in PFA at 4°C, washed with PBS, and transferred to a 30 % sucrose / PBS solution for 48 hours at 4°C. Brains were embedded in tissue freezing compound and 50 μm coronal sections were generated using a freezing microtome. Free floating thin sections were permeablised for 4-5 hours in block solution (PBS, 10 % horse serum, 0.5 % BSA, 0.5 % Triton X-100, 0.2 M glycine) then incubated with NeuN primary antibody diluted in block solution (1:1000) overnight at 4 °C. Slices were washed 4-5 times in PBS for 2 hours then incubated for 3-4 hours with secondary antibody (anti-rabbit Alexa Fluor 568; 1:1000; Invitrogen; Cat #A10042) and NeuroTrace Green Fluorescent Nissl Stain (1:2000; Invitrogen; Cat #N21480) at room temperature. Slices were then washed 4-5 times in PBS for 2 hours and mounted onto glass slides using ProLong Gold Antifade Mountant (Invitrogen; Cat #P36930). Sections were imaged on a Leica SP8 upright confocal laser scanning microscope using a X10/NA 0.45 objective. The tile function within the Leica software was used to acquire overlapping images over the whole section followed by the merge image processing function to stitch the tiles together.

### Protein biochemistry

Cultured hippocampal neurons from *Dnm1*^+/+^ or *Dnm1*^+/R237W^ littermates at DIV 14 were lysed directly into SDS (sodium dodecylsulfate) sample buffer (67 mM Tris, pH 7.4, 2 mM EGTA, 9.3% glycerol, 12% β-mercaptoethanol, bromophenol blue, 67 mM SDS) and boiled at 95°C for 10 minutes prior to Western blotting. Whole brain lysates were prepared from the brains of 3 week old and 6 week old age-matched *Dnm1*^+/+^ or *Dnm1*^+/R237W^ mice. Brain homogenates were prepared in RIPA buffer (10 mM Tris-HCl, pH 8.0, 1mM EDTA, 0.5 mM EGTA, 15 Triton X-100, 0.1% sodium deoxycholate, 0.1% SDS, 140 mM NaCl and 1 mM PMSF) and centrifuged in a Beckman-Coulter Optima-Max ultracentrifuge at 116,444 *g* for 40 min at 4°C. Protein concentration was determined using a Bradford (Applichem, Germany; A6932) assay following manufacturer’s instructions. SDS sample buffer was added to the lysates and samples were boiled for 10 min before loading on SDS-PAGE and blotting onto nitrocellulose membranes. Membranes were incubated with primary antibodies overnight at 4°C (Goat anti-amphyphysin-1 (Santa Cruz sc-8536) 1:500; rabbit anti-Eps15 (Santa Cruz sc-534) 1:1000; Goat anti-dynamin-1 (Santa Cruz sc-6402) 1:1000; rabbit anti-SNX9 (Santa Cruz sc-49143) 1:500; mouse anti-synaptotagmin-1 (Abcam ab13259) 1:500; rabbit anti-syndapin-1 (Abcam ab137390) 1:4000; goat anti-endophilin-A1 (Santa Cruz sc-10874) 1:1000; rabbit anti-C-src (Santa Cruz sc-19) 1:100; mouse anti-actin (Sigma Aldrich A4325) 1:50000). Secondary antibodies were incubated for 1h at room temperature (all Li-Cor, 1:10000; donkey anti-goat (red, 926-68074); donkey anti-goat (green, 926-32214); donkey anti-rabbit (green, 926-32213); donkey anti-mouse (green, 926-32212); goat anti-mouse (red, 926-68070). Membranes were imaged on an Odyssey 9120 Infrared Imaging System (LI-COR Biosciences) using *LI-COR* Image Studio Lite software (version 5.2) and analysed using ImageJ. The integrated density of signals was measured in rectangular ROIs of an identical size set around the protein expression bands.

### Mass Spectrometry

Synaptosomes were prepared from two-month-old *Dnm1*^+/+^ or *Dnm1*^+/R237W^ male mice as described^28^. Synaptosome pellets were dissolved in Urea lysis buffer (8M Urea in 50mM Tris-Cl and 1% sodium deoxycholate) and were quantified using the BCA method. 20 μg of total protein was used for proteomic sample preparation by suspension trapping (S-Trap) ^46^, as recommended by the supplier (ProtiFi, Huntington NY, USA). Samples were reduced with 5 mM Tris (2-carboxyethyl)phosphine (Pierce) for 30 min at 37°C, and subsequently alkylated with 5 mM IAM (Iodoacetamide) for 30 min at 37°C in the dark. After acidification with phosphoric acid, sample was cleaned and digested using Trypsin (1:20) as mentioned by in the manufacturer’s protocol using S-trap filter for 2 hours at 47°C and the digested peptides are eluted using 0.2% Formic acid and 50% Acetonitrile:0.2% formic acid. The eluted digested peptides were dried in speed vac and stored at -80°C.

The peptides were reconstituted in 30 µL of 0.1% formic acid and vortexed and 5 µL of each sample was injected on the mass spectrometer. Peptides were analysed by nanoflow-LC-MS/MS using a Orbitrap Q-Exactive-HF™ Mass Spectrometer (Thermo Scientific ^TM^) coupled to a Dionex™ Ultimate™ 3000. Samples were injected on a 100 μm ID × 5mm trap (Thermo Trap Cartridge 5mm) and separated on a 75 μm × 50 cm nano LC column (EASY-Spray™ LC Columns #ES803). All solvents used were HPLC or LC-MS Grade (Millipore™). Peptides were loaded for 5 minutes at 10 μL/min using 0.1% FA, 2% Acetonitrile in Water. The column was conditioned using 100% Buffer A (0.1% FA, 3% DMSO in Water) and the separation was performed on a linear gradient from 0 to 35% Buffer B (0.1% FA, 3% DMSO, 20% Water in Acetonitrile), over 140 minutes at 250 nL/min. The column was then washed with 90% Buffer B for 5 minutes and equilibrated 10 minutes with 100% Buffer A in preparation for the next analysis. Full MS scans were acquired from 350 to 1500 m/z at resolution 60,000 at m/z 200, with a target AGC of 3×10^6^ and a maximum injection time of 50 ms. MS/MS scans were acquired in HCD mode with a normalized collision energy of 25 and resolution 15000 using a Top 20 method, with a target AGC of 2×10^5^ and a maximum injection time of 50 ms. The MS/MS triggering threshold was set at 5E3 and the dynamic exclusion of previously acquired precursor was enabled for 45 s for DDA (Data-Dependent Acquisition) mode. For DIA (Data Independent Acquisition) mode the scan range was 385 to 1015 m/z, where MS/MS data was acquired in 24 m/z isolation windows at a resolution of 30,000.

Pooled peptides from all samples were fractionated on a Basic Reverse Phase column (Gemini C18, 3um particle size, 110A pore, 3 mm internal diameter, 250 mm length, Phenomenex #00G-4439-Y0) on a Dionex Ultimate 3000 Off-line LC system. All solvent used were HPLC grade (Fluka). Peptides were loaded on column for 1 minute at 250 μL/min using 99% Buffer A (20mM Ammonium Formate, pH=8) and eluted for 48 minutes on a linear gradient from 2 to 50% Buffer B (100% ACN). The column is then washed with 90% Buffer B for 5 minutes and equilibrated for 5 minutes for the next injection. Peptide elution was monitored by UV detection using at 214 nm. Fractions were collected every 45 s from 2 min to 60 min for a total of 12 fractions. Non-consecutive concatenation of every 13th fraction was used to obtain 12 pooled fractions (Pooled Fraction 1: Fraction 1 + 13 + 25 + 37, Pooled Fraction 2 : Fraction 2 + 14 + 26 + 38 …).

### Data Analysis

Label-free quantitative analysis was performed using the data set acquired in DIA mode. Peptide identification was carried out using a library generated using both DDA and DIA datasets using Spectronaut^TM^ version 15.0. The library was generated using the Pulsar algorithm integrated in Spectronaut using Mus musculus FASTA using 1% FDR. The maximum of missed cleavage was set to 2 using Trypsin/P enzyme. Carbamidomethylation (C) was set as fixed modification and acetylation (Protein N term), oxidation (M), deamination (NQ), were set as variable modifications. The library consisted spectra information of 5906 proteins in total. DIA data set for both WT and HET was searched using this library quantified 4237 proteins in total. Statistical analysis was done using R script and limma package was used for making contrasts.

### Analysis of mass spectrometry data

Gene Ontology (GO) terms enrichment analysis on the upregulated and downregulated proteins was performed against Mus musculus background using Database for Annotation, Visualization and Integrated Discovery (DAVID). Detailed enrichment analysis are available in **Extended Data Table 3**. Network analysis of the upregulated and downregulated proteins were performed using the STRING web tool (v.11.5).

Enrichment analysis of the full protein list were performed using ShinyGO v0.76.2 for the Cellular Component and Biological Pathways, selected by FDR and sorted by FoldEnrichment and using the synapse specific database SynGO ^47^ against the “brain expressed” background, setting medium stringency and second level terms as labels for Cellular Component representation and top levels terms as labels for Biological Pathways representation (**Extended Data Figure 3**).

### GTPase assays

A colorimetric assay was used to quantify GTPase activity of the different mCer-Dyn1 mutants ^48^. HEK293T cells transfected with mCer-Dyn1 plasmids were harvested 48 h after transfection with a 1:1 ratio of Lipofectamine2000 to plasmid. The cells were resuspended in 1 ml of sucrose lysis buffer (250 mM sucrose, 3 mM imidazole pH 7.4 supplemented with 2 μl/ml protease inhibitors and 1 mM phenylmethane sulfonyl fluoride) and mechanically broken using a primed ball-bearing cell cracker (EMBL, Heidelberg, Germany). Anti-GFP VHH coupled to agarose beads for immunoprecipitation of GFP-fusion proteins (GFP-Trap; ChromoTek GmbH, Germany; gta-20) was used for immunoprecipitation of mCerN1, mCer-Dyn1WT or mutant mCer-Dyn1 according to manufacturer’s instructions. A Bradford (Applichem, Germany; A6932) assay was performed according to manufacturer’s instructions to determine protein concentration of GFP-Trap-bound mCerN1 or mCer-Dyn1. GFP-Trap-bound mCer-Dyn1 mutants were diluted to a concentration of 1 μM in GTPase assay buffer (20 mM HEPES pH 7.5, 50 mM KCl-this low salt concentration allows for the oligomerisation of dynamin ^49^, 2 mM MgCl_2_). For each reaction, 20 μl of 2 mM GTP diluted in GTPase assay buffer and 20 μl of 1 μM stock of mCer-Dyn1 was incubated for 30 min at 37°C after which 0.5 M EDTA pH 8.0 was added to terminate the reaction. 300 ul of filtered Malachite green solution (34 mg Malachite green carbinol base dissolved in 40 mL of 1 N HCl added to 1 g of ammonium molybdate tetrahydrate diluted in 14 mL of 4 N HCl up to 100 mL with ddH_2_O) was added to each reaction. The change in colour of malachite green was quantified using a plate reader to measure the absorbance at 650 nm. The amount of inorganic phosphate released was calculated using the standard curve.

### Acute slice preparation

Horizontal hippocampal slices (350 µm) were prepared from *Dnm1*^+/R237W^ and *Dnm1*^+/+^ littermate control mice (P19-25 of either sex). Excised brains were rapidly transferred to chilled (2 – 5°C) carbogenated sucrose-modified artificial cerebrospinal fluid (saCSF in mM: NaCl 86, NaH_2_PO_4_ 1.2, KCl 2.5, NaHCO_3_ 25, glucose 25, sucrose 50, CaCl_2_ 0.5, and MgCl_2_ 7) for 2 minutes and subsequently sliced in the same solution using a vibrating microtome (Leica VT1200S). Slices were allowed to recover for 1 hour at 33°C in carbongenated standard aCSF which contained (mM): NaCl 126, KCl 3, NaH_2_PO_4_ 1.2, NaHCO_3_ 25, glucose 15, CaCl_2_ 2, and MgCl_2_ 2.

### Electrophysiology

For recording, slices were transferred to an immersion chamber continuously perfused with standard aCSF (MgCl_2_ 1 mM) maintained at 32°C using an in-line Peltier heater (Scientifica, Uckfield, UK). A cut was made between CA2 and CA1 (identified as the medial termination of stratum lucidum) to ablate recurrent activity. Whole-cell patch-clamp recordings were made from visually identified pyramidal neurons in the CA1 region using pulled borosilciate electrodes (4-7 MΩ). The intracellular solution for evoked and intrinsic properties experiments consisted of (mM): K-gluconate 142, KCl 4, EGTA 0.5, HEPES 10, MgCl_2_ 2, Na_2_ATP 2, Na_2_GTP 0.3, and Na_2_-creatine 10. For miniature excitatory post-synaptic current (mEPSC) recordings, a caesium-based intracellular solution was used (mM): Cs-gluconate 140, CsCl 3, EGTA 0.2, HEPES 10, QX-314 chloride 5, MgATP 2, NaATP 2, Na_2_GTP 0.3, and phosphocreatine 10. Excitatory currents were recorded in the presence of picrotoxin (50 µM) with cells voltage-clamped at -70 mV, with the further addition of TTX (300 nM) for mEPSC recording. For experiments with BMS-204352, the drug was dissolved in DMSO and TWEEN® 80 before adding to standard aCSF. Final drug concentration was 30 μM, with the vehicles both at 0.03% v/v.

Recording protocols: Intrinsic properties were recorded in current-clamp mode. All other recordings were made under voltage-clamp. Currents were low pass filtered at 3–10 kHz and sampled at 10-20 kHz, using Clampex 10 softwarMe (pClamp 10, Molecular Devices, San Jose, USA). For evoked recordings, Schaffer collaterals were stimulated with a patch electrode (∼1–2 MΩ) filled with aCSF and positioned in stratum radiatum, connected to an isolated constant current stimulator (Digitimer, Welwyn Garden City, UK). In all cases, the stimulus intensity was set to evoke a current of ∼200 pA following a 50 µs pulse. Stimulus was delivered at either: paired pulses (interval 10 – 500 ms, pairs 30 s apart), or long trains (either 10 or 40 Hz for 15 s, four repeats delivered 4 minutes apart). Data were analysed offline using either the open source Stimfit software package (intrinsic properties) or Clampfit from the pClamp 10 software suite (all EPSCs). Cells were excluded from analysis if series resistance varied by more than 20% during recording.

RRP size and its replenishment were determined using approaches described in ^21^. Briefly RRP was calculated by plotting the cumulative EPSC amplitude from 40 Hz 15 s trains, and performing a linear regression on the last 1 s of that plot. The y-intercept of this regression line denotes RRP size (**Figure 4g**). Replenishment rate is represented by the slope of the regression line. Pr was calculated as amplitude of the first evoked EPSC divided by the effective RRP size ^50^.

### Surgery for *in vivo* electrophysiology

*Dnm1*^+/R237W^ and *Dnm1*^+/+^ littermate control mice of either sex aged 8 weeks were anaesthetized with isoflurane and mounted on a stereotaxic frame (David Kopf Instruments, USA). Pairs of local LFP electrodes (Ø = 50.8 µm, Teflon insulated stainless steel, A-M Systems, USA) were implanted targeting dorsal hippocampus bilaterally (1.85 mm caudal, 1.25 mm lateral from bregma and 1.40 mm ventral from brain surface), ventral hippocampus bilaterally (3.3 mm caudal, 3.3 mm lateral from bregma and 2.9 mm ventral from the brain surface), left motor cortex (1.55 mm caudal, 1.88 mm left from bregma and on the brain surface), right somatosensory cortex (1.3 mm caudal, 2.0 mm lateral from bregma and on the brain surface) and the midline cerebellum (5.7 mm caudal, 0 mm lateral from bregma and on the brain surface). Two miniature ground screws (Yahata Neji, M1 Pan Head Stainless Steel Cross, RS Components, Northants, UK) were attached over the cerebellum (5.0 mm caudal, 2 mm lateral) to serve as ground as well as three additional screws for structural support. The electrodes were attached to an electronic interface board (EIB-16, Neuralynx, USA). The electrode assemblies were fixed to the skull using a combination of UV activated cement (3M Relyx Unicem 2 Automix, Henry Schein, Gillingham, UK) and dental cement (Simplex Rapid, Kemdent, Swindon, UK).

### *In vivo* LFP recordings

Mice were placed in 50 x 50 cm square arenas and connected for recordings to an RHD 16-channel recording headstage (Intantech, USA) through an electrical commutator (Adafruit, USA) and an acquisition board (OpenEphys, USA). LFP signals were sampled at 1 kHz and referenced to ground. Mice were video-recorded during stimulation sessions at 9.98 frames/s (C270 HD webcam, Logitech, USA). A 1 s light pulse from a blue LED (blue = 465 nm, Plexon, USA) mounted on each commutator was triggered by a Master-8 (AMPI) every five min to synchronize jump timestamps in video and LFP recordings.

### Behavioural experiments

For the open field assay, 6- to 8-week-old *Dnm1*^+/R237W^ and *Dnm1*^+/+^ littermate controls of either sex were placed in an open field arena 50 cm x 50 cm for 30 mins for 5 consecutive days. The first day in the arena served as habituation. On days 2 (test 1) and 4 (test 2) mice received 2 mg/g BMS-204352 or vehicle (DMSO 1/80; Tween 80 1/80; 0.9% NaCl) administered by intraperitoneal injection (as described in ^24^) in a counterbalanced manner as described in **Figure 8a**. No injections were administered on days 3 and 5 (washout). Injections were administered 20 min prior to start of experiment to ensure maximal brain BMS-204352 concentration for duration of time in open field arena. Activity was recorded at 9.89 fps from both the top view and side view of the arena using Logitech cameras (C270 HD webcam, Logitech) with up to 4 animals being recorded simultaneously in individual arenas. Jumping behaviour was scored using Behavioral Observation Research Interactive Software (BORIS, University of Torino ^51^) which allowed for each jump to be logged in time.

### Analysis of mouse movement and location in behavioural tasks

DeepLabCut (DLC) was used to compare the movement and position of mice ^52^. The tail base was used for analysis, since it provided the most accurate approximation of movement and position in two-dimensions. DLC tracked the movement of the animal for the duration of each video and provided an output for its X- and Y-coordinates at every frame. A loop was used to iterate the predicted tail base coordinates in each video and calculate the distance an animal travelled between each frame. These distances were summed across frames to determine total distance covered in the 30 minute experiment. Videos were then grouped based on their camera angle. For each angle, DLC was used to assign coordinates to the corners of the animal’s arena, which allowed conversion of DLC units into centimetres. To determine the time an animal spent in the centre and along the walls of the arena, different camera angles were used. For each angle, the dimensions of the animal’s arena were approximated to create an “outer” and an “inner” box. Each box contained half the total area of the arena. The number of tail base coordinates found within both boxes were totalled (time spent in centre), as well as those only found within the large box (time spent at edges).

### Statistical analysis

Experimenters were blinded to the genotype of both animals and cells for all experiments and data analysis. Statistical analysis was performed using GraphPad Prism 8.4.3. Statistical analysis for paired behaviour data was analysed using IBM SPSS Statistics v29. No statistical methods were used to predetermine sample sizes and no randomization procedures were applied. Statistical tests were applied based on the distribution of the datasets measured using D’Agostino-Pearson normality test. Significance was set at ns P > 0.05, ^∗^P < 0.05, ^∗∗^P < 0.01, ^∗∗∗^P< 0.001, ^∗∗∗∗^ P < 0.0001. Mann-Whitney (two-tailed), Wilcoxon matched-pairs signed rank (two-tailed), and Kruskal-Wallis with Dunns *post-hoc* tests were used to compare non-Gaussian data sets. Student’s t test (two-tailed) and analyses of variance (ANOVA) followed by Dunnett’s *post-hoc* test were used to compare normally distributed data sets. General linear model (repeated measures) was used to determine genotype effects, treatment effects and interactions. Bonferroni multiple comparisons test was used for multi-group comparisons where appropriate. Information about sample sizes, statistical tests used to calculate P values and the numeric values of the results are specified in figure legends and **Extended Data Table 4**.

