## Supplementary Data for "Reversal of cell, circuit and seizure phenotypes in a mouse model of *DNM1* epileptic encephalopathy"

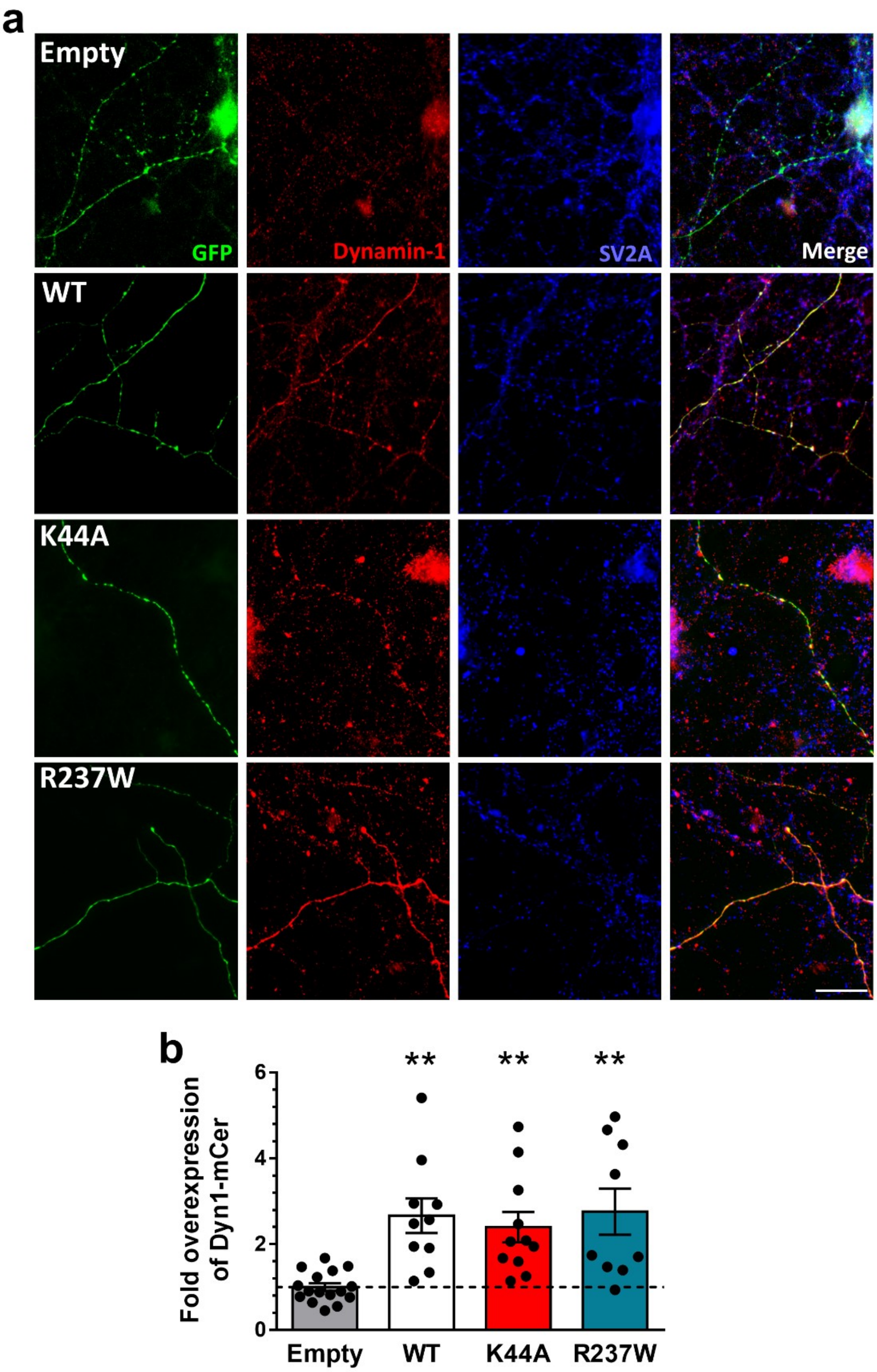

**Figure S1** – *Expression levels of Dyn1$_{WT}$-mCer plasmids.* Primary cultures of hippocampal neurons were transfected with synaptophysin-pHluorin (sypHy) and either mCer (Empty), Dyn1$_{WT}$-mCer, Dyn1$_{K44A}$-mCer or Dyn1$_{R237W}$-mCer between 11-13 DIV. At 13-15 DIV, cultures were fixed and immunostained for the presence of mCer (GFP), dynamin-1 and SV2A. (**a**) Representative images, scale bar = 20 μm. (**b**) Average fold over-expression of Dyn1-mCer plasmids, normalised to endogenous dynamin-1 levels ± SEM. One-way ANOVA, n=16 Empty, n=10 Dyn1$_{WT}$-mCer, n=11 Dyn1$_{K44A}$-mCer, n=9 Dyn1$_{R237W}$-mCer, All against Empty ** p=0.0015 Dyn1$_{WT}$-mCer, ** p=0.0065 Dyn1$_{K44A}$-mCer, ** p=0.0012 Dyn1$_{R237W}$-mCer.

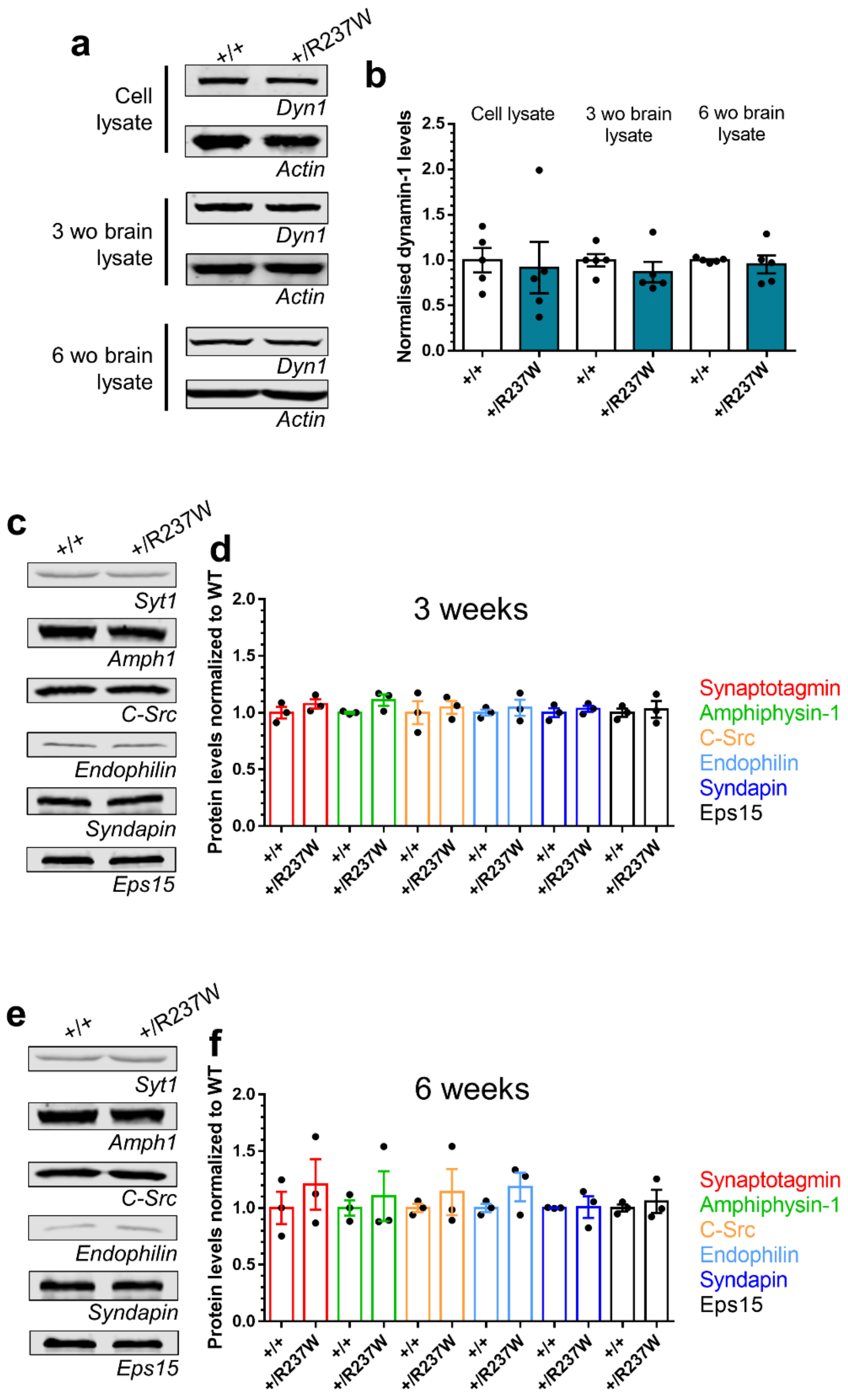

**Figure S2** – *Dynamin-1 and other synaptic protein levels in Dnm1*$^{+/R237W}$ *mice.* Lysates were generated form either primary cultures of either *Dnm1*$^{+/+}$ or *Dnm1*$^{+/R237W}$ hippocampal neurons or from the brains of either 3 week or 6 week old *Dnm1*$^{+/+}$ or *Dnm1*$^{+/R237W}$ mice. (**a**) Representative blots for dynamin-1 levels in *Dnm1*$^{+/+}$ or *Dnm1*$^{+/R237W}$ lysates. (**b**) Average levels of dynamin-1 normalised to *Dnm1*$^{+/+}$ lysates ± SEM (Unpaired t test with Welch's correction, all n=5 samples, p=0.801 cell lysate, p=0.359 3 month, p=0.671 6 month). (**c,e**) Representative blots display levels of Synaptotagmin-1 (Syt1), Amphiphysin-1 (Amph1), C-src, Endophilin, Syndapin and Eps15 in either 3 week (**c**) or 6 week (**e**) lysates. (**d,f**) Quantification of protein levels normalised to *Dnm1*$^{+/+}$ ± SEM in either 3 week (**d**) or 6 week (**f**) lysates (n=3 independent brain lysate preparations for all, all ns, Mann-Whitney test).

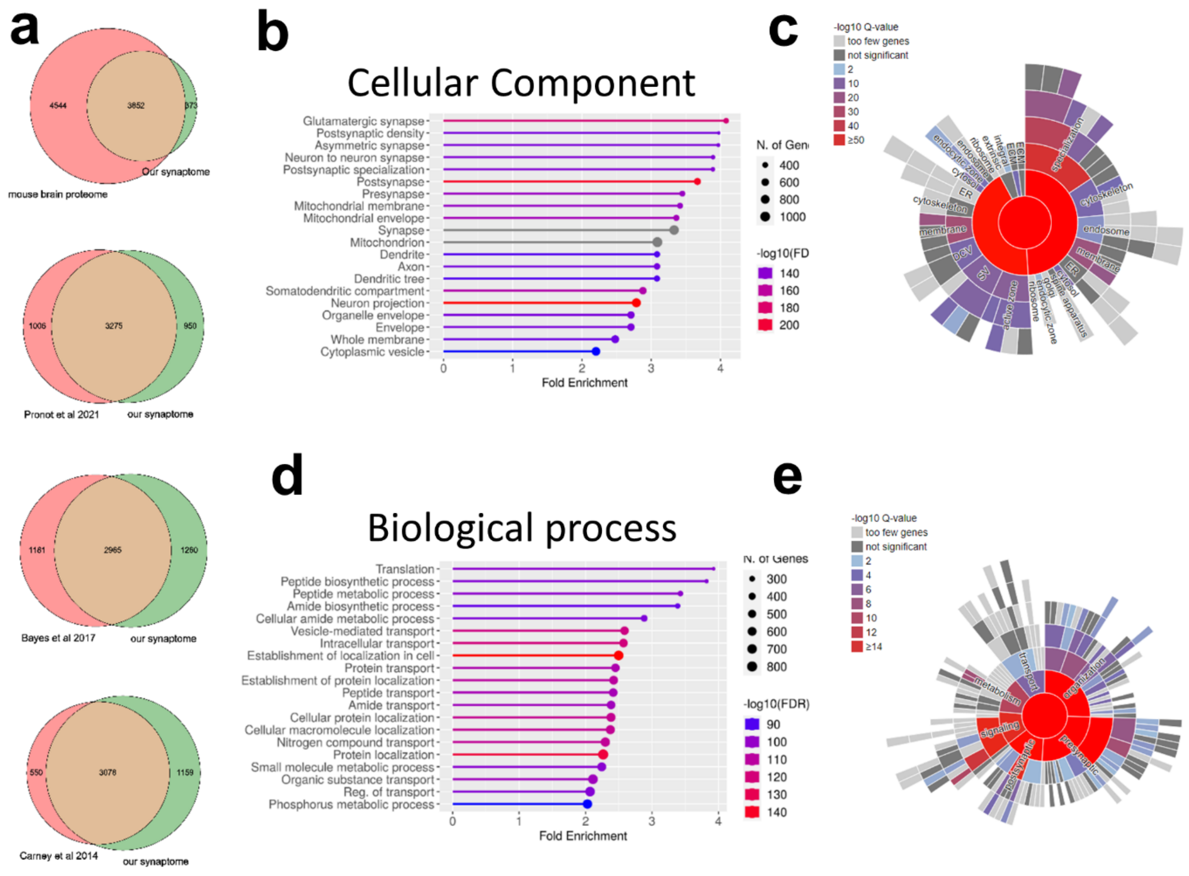

**Figure S3** – *Proteomic data.* (**a**) Venn diagrams showing overlap between the synaptome of this study and other published synaptosome proteomes. Terms enrichment analysis of 4237 synaptic proteins found by MS for GO Cellular Components (**b**) and Biological Process (**d**) using ShinyGO databases and for the Cellular Components (**c**) and Biological Pathways (**e**) using SynGO synaptic component curator tool.

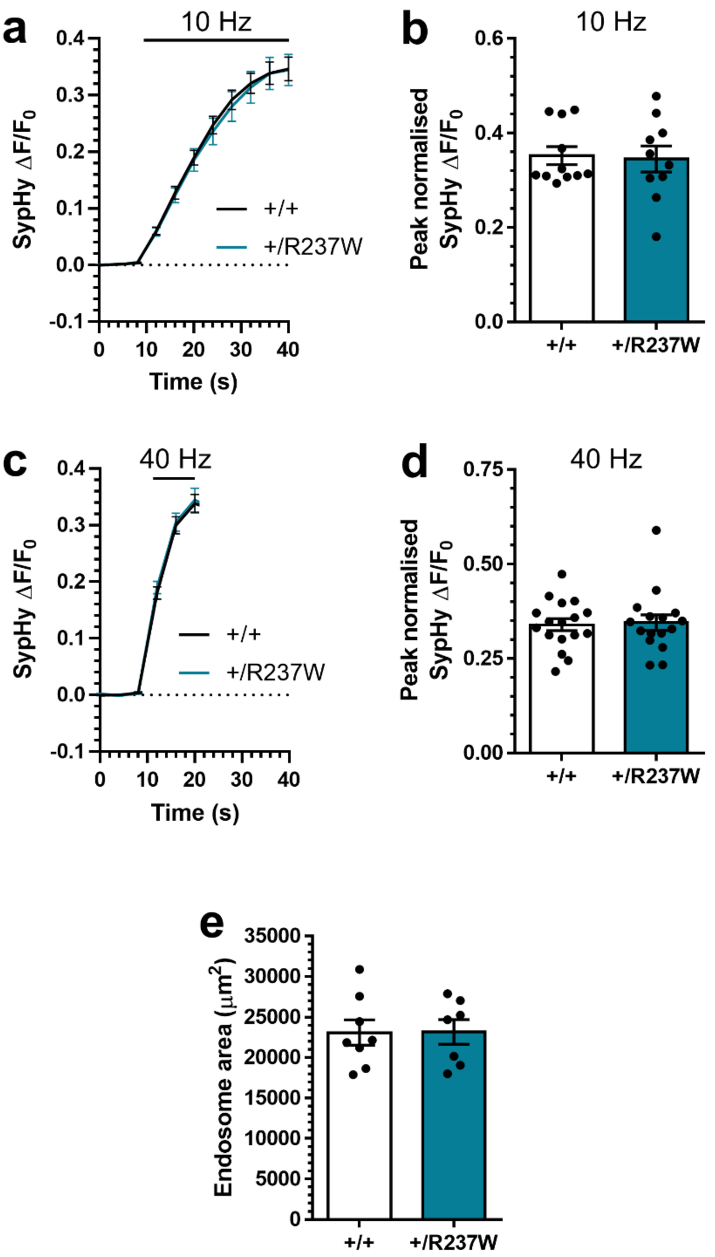

**Figure S4** – *SV pools and endosomes in Dnm1$^{+/R237W}$ mice.* (**a-d**) Primary cultures of hippocampal neurons prepared from either *Dnm1*$^{+/+}$ and *Dnm1*$^{+/R237W}$ embryos were transfected with synaptophysin-pHluorin (sypHy) between 7-9 DIV. At 13-15 DIV, cultures were stimulated with a train of either (**a,b**) 300 action potentials (10 Hz) or (**c,d**) 400 action potentials (40 Hz) in the presence of 1 μM bafilomycin-A1. Immediately after stimulation, cultures were pulsed with $NH_4Cl$ imaging buffer. (**a,c**) Average sypHy response ($\Delta F/F_0$ ± SEM) to either 10 Hz (**a**) or 40 Hz (**c**) stimulation, normalised to the $NH_4Cl$ challenge peak. (**b,d**) Peak level of sypHy fluorescence ($\Delta F/F_0$ ± SEM) normalised to the $NH_4Cl$ challenge (**b** Unpaired t test, n=11 *Dnm1*$^{+/+}$, n=10 *Dnm1*$^{+/R237W}$, p=0.839; **d** Mann-Whitney test, n=17 *Dnm1*$^{+/+}$, n-16 *Dnm1*$^{+/R237W}$, p=0.919). (**e**) *Dnm1*$^{+/+}$ and *Dnm1*$^{+/R237W}$ neurons were stimulated with a train of 400 action potentials (40 Hz) in the presence of 10 mg/ml HRP. Average number HRP-labelled endosome diameter is displayed ± SEM (Unpaired t test, n=8 *Dnm1*$^{+/+}$, n=7 *Dnm1*$^{+/R237W}$ p=0.974).

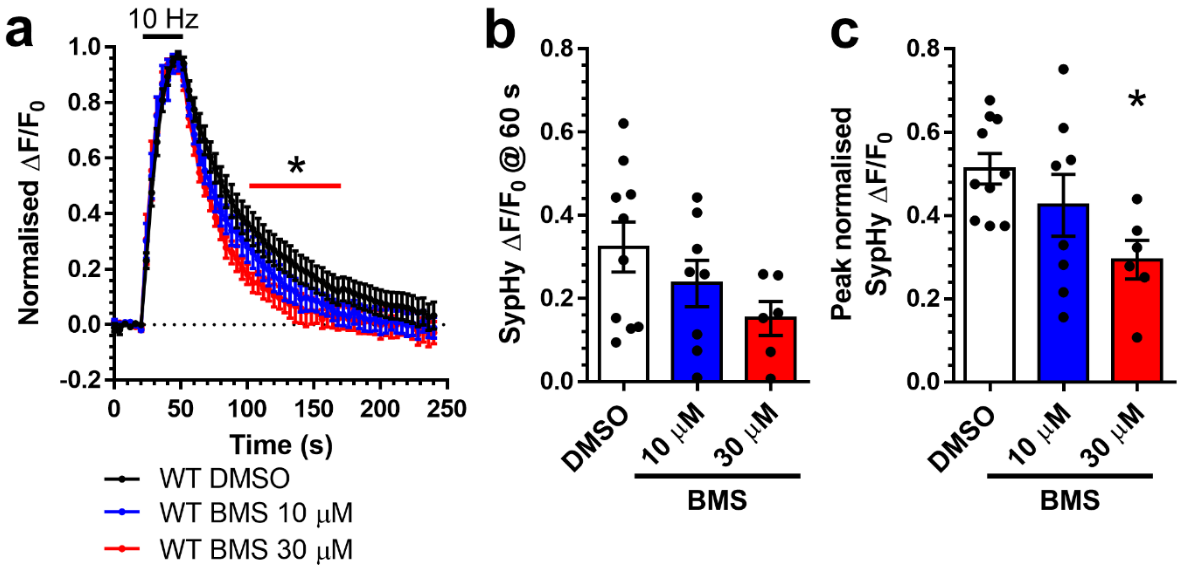

**Figure S5** – *BMS-204352 accelerates SV endocytosis.* Primary cultures of hippocampal neurons prepared from *Dnm1*$^{+/+}$ embryos were transfected with synaptophysin-pHluorin (sypHy) and Dyn1$_{WT}$-mCer between 11-13 DIV. At 13-15 DIV, cultures were stimulated with a train of 300 action potentials (10 Hz) in the presence of either 10 μM or 30 μM BMS-204352 or a vehicle control (DMSO). Cultures were pulsed with $NH_4Cl$ imaging buffer 180 s after stimulation. (**a**) Average sypHy response ($\Delta F/F_0 \pm$ SEM) normalised to the stimulation peak. Two-way ANOVA *$p< 0.05$. (**b**) Average level of sypHy fluorescence ($\Delta F/F_0 \pm$ SEM) at 60 s (One-way ANOVA, n=10 DMSO, N=8 10 μM, n=6 30 μM, all ns). (**c**) Peak level of sypHy fluorescence ($\Delta F/F_0 \pm$ SEM) normalised to the $NH_4Cl$ challenge (One-way ANOVA, n=10 DMSO, N=8 10 μM, n=6 30 μM, *p=0.0217 DMSO vs 30 μM).

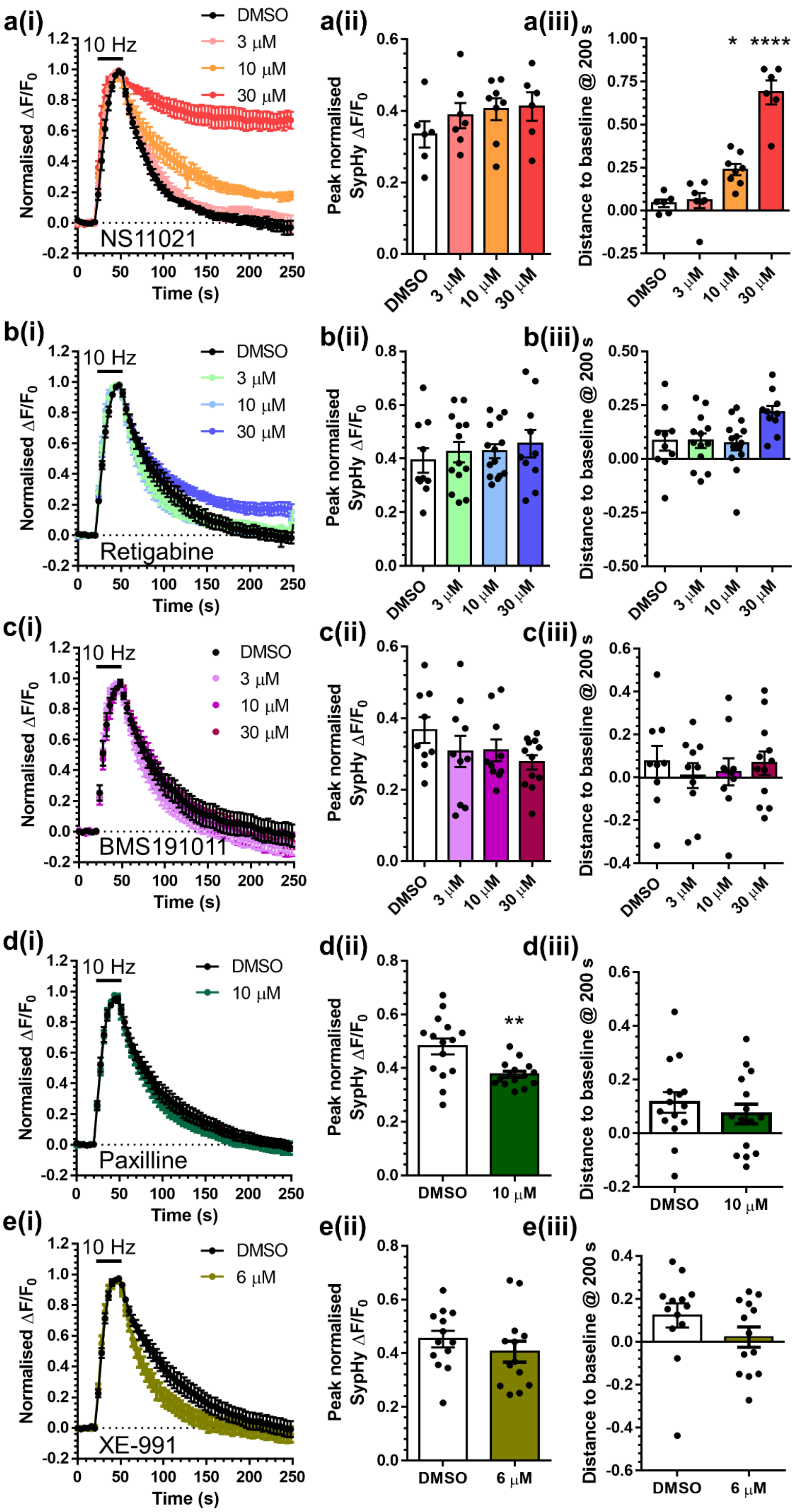
a(i)
10 Hz
DMSO
3 μM
10 μM
30 μM
NS11021
Normalised ΔF/F0
Time (s)
a(ii)
Peak normalised SypHy ΔF/F0
DMSO
3 μM
10 μM
30 μM
a(iii)
Distance to baseline @ 200 s
*
****
b(i)
10 Hz
Retigabine
b(ii)
b(iii)
c(i)
10 Hz
BMS191011
c(ii)
c(iii)
d(i)
10 Hz
Paxilline
d(ii)
**
d(iii)
e(i)
10 Hz
XE-991
6 μM
e(ii)
e(iii)

**Figure S6** – *Effect of potassium channel modulators on SV recycling.* Primary cultures of hippocampal neurons prepared from *Dnm1*$^{+/+}$ embryos were transfected with synaptophysin-pHluorin (sypHy) between 7-9 DIV. At 13-15 DIV, cultures were stimulated with a train of 300 action potentials (10 Hz) in the presence of 3 μM, 10 μM or 30 μM of either NS11021 (**a**), Retigabine (**b**) or BMS-911011 (**c**) or a vehicle control (DMSO). In identical separate experiments, cultures were stimulated in the presence of 10 μM Paxilline (**d**) or 6 μM XE-991 (**e**) or a vehicle control (DMSO). Cultures were pulsed with $NH_4Cl$ imaging buffer 180 s after stimulation. (**all i**) Average sypHy response ($\Delta F/F_0$ ± SEM) to 10 Hz stimulation, normalised to stimulation. (**all ii**) Average peak level of sypHy fluorescence ($\Delta F/F_0$ ± SEM) normalised to the $NH_4Cl$ challenge (NS11021, one-way ANOVA, n=6 DMSO, n=7 3 μM, n=8 10 μM, n=6 30 μM, all ns; Retigabine, one-way ANOVA, n=10 DMSO, n=13 3 μM, n=14 10 μM, n=10 30 μM, all ns; BMS-911011, one-way ANOVA, n=9 DMSO, n=10 3 μM, n=10 10 μM, n=12 30 μM, all ns; Paxilline, Unpaired t test n=15 both DMSO and 10 μM, p=0.003; XE-991, Unpaired t test n=13 both DMSO and 6 μM, p=0.361). (**all, iii**) Average level of sypHy fluorescence ($\Delta F/F_0$ ± SEM) at 200 s (NS11021, one-way ANOVA, n=6 DMSO, n=7 3 μM, n=8 10 μM, n=6 30 μM, *p=0.011 DMSO vs 10 μM, ****p<0.00001 DMSO vs 30 μM; Retigabine, one-way ANOVA, n=10 DMSO, n=13 3 μM, n=14 10 μM, n=10 30 μM, all ns; BMS-911011, one-way ANOVA, n=9 DMSO, n=10 3 μM, n=10 10 μM, n=12 30 μM, all ns; Paxilline, Unpaired t test n=15 both DMSO and 10 μM, p=0.427; XE-991, Mann-Whitney test n=13 both DMSO and 6 μM, p=0.152).

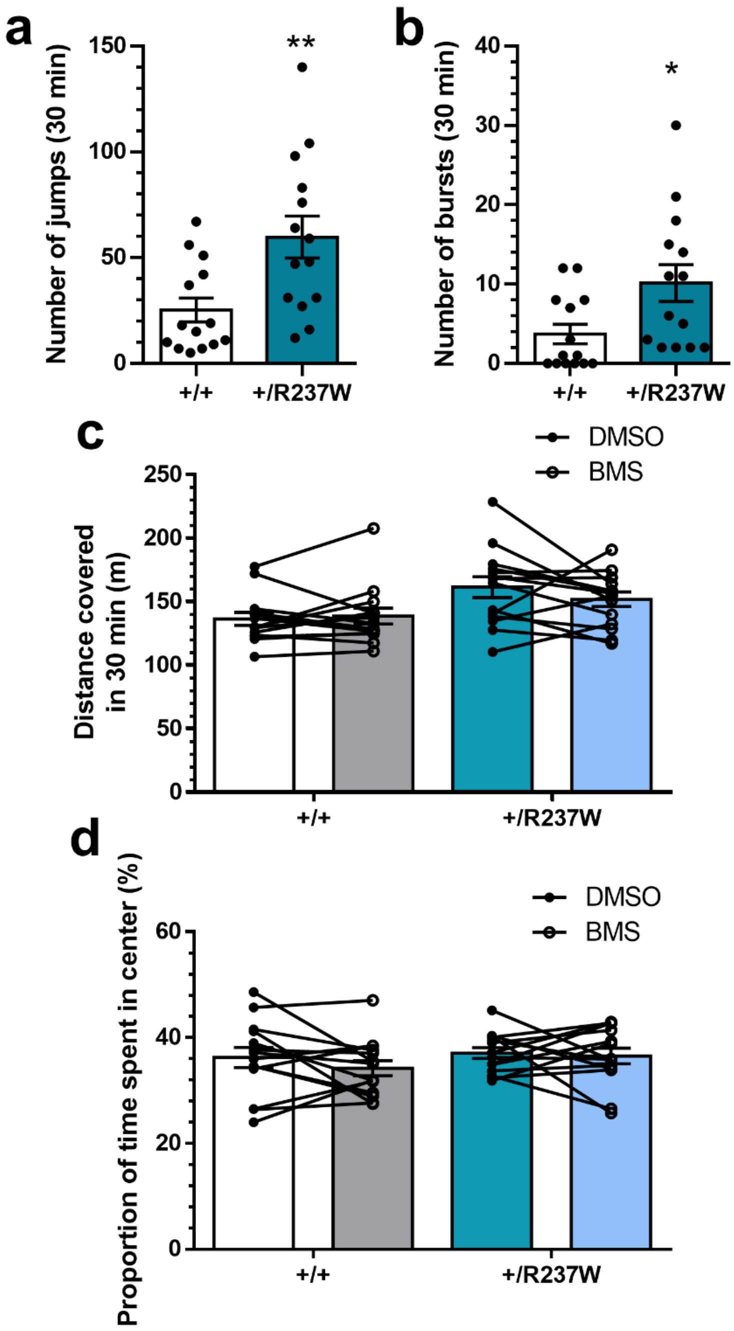

**Figure S7** – *Dnm1$^{+/R237W}$ mice display increased myoclonic jumping and jumping bursts in open field task. Dnm1$^{+/+}$* and *Dnm1$^{+/R237W}$* mice were placed in an open field chamber for a 30 min period for 5 days. After habituation on day 1, mice were dosed with either BMS-204352 or a vehicle control (DMSO) on days 2 and 4, and no treatment (washout) on days 3 and 5. Delivery of drug treatment was interleaved between days 2 and 4. (**a,b**) Comparison of *Dnm1$^{+/+}$* and *Dnm1$^{+/R237W}$* mice in the washout phase of day 5. Average number of myoclonic jumps (**a**), jumping bursts (**b**) is displayed ± SEM (Unpaired t test; n=14 for all; **a** ** p=0.0066; **b** * p=0.0238). (**c,d**) Comparison of *Dnm1$^{+/+}$* and *Dnm1$^{+/R237W}$* mice in the test phase (day 2 and 4). Average distance covered (**c**) and time spent in the centre of the area within the 30 min time period (**d**) is displayed ± SEM, General linear model (repeated measures) n=14 for all, all ns.

**Extended Data Table 1 –** Table displaying protein identified synaptosomes derived from *Dnm1*$^{+/+}$ and *Dnm1*$^{+/R237W}$ mice. Proteins significantly increased in *Dnm1*$^{+/R237W}$ synaptosomes are highlighted in orange (see column B for log2 ratio), whereas those decreased are highlighted in blue. Dynamin-1 is highlighted in green.

| | Standard aCSF | |
|---|---|---|
| | **WT** | **Het** |
| **Resting membrane potential (mV)** | -66.38 | -65.20 |
| **Baseline (mV)** | -70.76 | -70.48 |
| **Step with 10 pA injection (mV)** | -1.305 | -1.332 |
| **Input resistance (MΩ)** | 130.5 | 133.2 |
| **Membrane decay time τ (ms)** | 18.26 | 21.17 |
| **Capacitance (MF)** | 139.3 | 160.8 |
| **Rheobase (pA)** | 107.9 | 103.9 |
| **AP threshold (mV)** | -43.22 | -44.47 |
| **AP peak (mV)** | 82.37 | 85.29 |
| **AP rise time 20 to 80% (ms)** | 0.1593 | 0.1765 |
| **AP width at half peak height (ms)** | 0.873 | 0.976 |
| **Max rise rate (mV/ms)** | 401.6 | 370.9 |
| **Max decay rate (mV/ms)** | 95.77 | 86.47 |
| **Rise/Decay** | 4.14 | 4.24 |
| **Sag current as % of steady state** | 24.9 | 28.7 |
| **Rebound potential (mV)** | 4.01 | 4.81 |
| **mAHP (mV)** | -8.83 | -8.35 |
| **fAHP (mV)** | -4.78 | -4.09 |
| **Max firing frequency (Hz)** | 50.7 | 49.4 |

**Extended Data Table 2** – *Intrinsic properties of Dnm1*$^{+/R237W}$ *CA1 neurons*. Acute hippocampal slices from P19-25 *Dnm1*$^{+/+}$ or *Dnm1*$^{+/R237W}$ mice were prepared and the intrinsic properties of CA1 neurons were determined using whole-cell patch clamp recording. The parameters monitored are outlined above. Highlighted pairs = $p < 0.05$ unpaired two-tailed t test.

**Extended Data Table 3 –** DAVID analysis of up and down-regulated proteins in *Dnm1*$^{+/R237W}$ synaptosomes.

| Figure 1 | Group | Mean±SEM | n = # of experiments/coverslips N = # of neuronal preparations | Comparison | P | Statistical test |
|---|---|---|---|---|---|---|
| **Fig1a** | Empty | 62.6 ± 1.1 μM | 4 | WT vs Empty | <0.0001 | One-way ANOVA with Dunnett's multiple comparison test |
| | WT | 119.6 ± 5.8 μM | 4 | | | |
| | K44A | 93.6 ± 7.2 μM | 4 | WT vs K44A | 0.0138 | |
| **Fig1b** | Empty | 63.8 ± 3.5 μM | 3 | WT vs Empty | <0.0001 | |
| | WT | 123.2 ± 2.7 μM | 3 | | | |
| | R237W | 102.8 ± 3.9 μM | 3 | WT vs R237W | 0.0093 | |
| **Fig1d** | mCer | 0.12 ± 0.034 | 9/4 | WT vs mCer | 0.978 | |
| | WT | 0.11 ± 0.039 | 10/4 | | | |
| | K44A | 0.31 ± 0.044 | 17/4 | WT vs K44A | 0.0046 | |
| **Fig1e** | mCer | 0.44 ± 0.036 | 9/4 | WT vs mCer | 0.086 | |
| | WT | 0.55 ± 0.044 | 10/4 | WT vs K44A | 0.472 | |
| | K44A | 0.50 ± 0.028 | 17/4 | | | |
| **Fig1g** | WT | 0.17 ± 0.047 | 16/5 | WT vs R237W | 0.0007 | Unpaired t test |
| | R237W | 0.50 ± 0.081 | 8/5 | | | |
| **Fig1h** | WT | 0.47 ± 0.042 | 16/5 | WT vs R237W | 0.480 | |
| | R237W | 0.43 ± 0.042 | 8/5 | | | |

| Figure 2 | Group | Mean±SEM | n = # of neuronal preparations | Comparison | P | Statistical test |
|---|---|---|---|---|---|---|
| **Fig2c** | WT Syt1 | 1.00 ± 0.053 | 3 | WT vs R237W Syt1 | 0.900 | Mann-Whitney test |
| | R237W Syt1 | 0.93 ± 0.12 | 3 | | | |
| | WT Amph1 | 1.00 ± 0.076 | 3 | WT vs R237W Amph1 | 0.700 | |
| | R237W Amph1 | 0.91 ± 0.13 | 3 | | | |
| | WT C-Src | 1.00 ± 0.053 | 3 | WT vs R237W C-Src | 0.700 | |
| | R237W C-Src | 0.88 ± 0.17 | 3 | | | |
| | WT Endo | 1.00 ± 0.053 | 3 | WT vs R237W Endo | 0.400 | |
| | R237W Endo | 0.83 ± 0.12 | 3 | | | |
| | WT Syd | 1.00 ± 0.053 | 3 | WT vs R237W Syd | 0.900 | |
| | R237W Syd | 0.93 ± 0.12 | 3 | | | |
| | WT Eps15 | 1.00 ± 0.053 | 3 | WT vs R237W Eps15 | 0.900 | |
| | R237W Eps15 | 0.90 ± 0.16 | 3 | | | |

| Figure 3 | Group | Mean±SEM | n = # of coverslips/ N = # of neuronal preparations | Comparison | P | Statistical test |
|---|---|---|---|---|---|---|
| **Fig3d** | WT | 0.12 ± 0.042 | 9/4 | WT vs R237W | 0.045 | Mann-Whitney test |
| | R237W | 0.29 ± 0.052 | 12/4 | | | |
| **Fig3e** | WT | 0.33 ± 0.023 | 9/4 | WT vs R237W | 0.237 | Unpaired t test |
| | R237W | 0.41 ± 0.031 | 12/4 | | | |
| **Fig 3g** | WT | 0.13 ± 0.029 | 9/4 | WT vs R237W | 0.003 | |
| | R237W | 0.30 ± 0.037 | 11/4 | | | |
| **Fig 3h** | WT | 0.44 ± 0.0.38 | 9/4 | WT vs R237W | 0.751 | |
| | R237W | 0.42 ± 0.049 | 11/4 | | | |
| **Fig 3k** | WT | 3.34 ± 0.160 | 8/4 | WT vs R237W | 0.0393 | |
| | R237W | 2.90 ± 0.099 | 7/4 | | | |

| | | | | | | |
|---|---|---|---|---|---|---|
| Figure 4 | Group | Mean±SEM | n = # slices / N = # of animals | Comparison | P | Statistical test |
| **Fig 4b** | WT | 2.09 ± 0.28 | 10/6 | WT vs R237W | 0.0008 | Mann-Whitney test |
| | R237W | 0.92 ± 0.13 | 12/7 | | | |
| **Fig 4c** | WT | -25.2 ± 2.32 | 10/6 | WT vs R237W | 0.665 | |
| | R237W | -25.6 ± 1.50 | 12/7 | | | |
| **Fig 4d** | WT | | 11/5 | WT vs R237W Overall | <0.0001 | Two-way ANOVA with Fisher's LSD |
| | R237W | | 10/6 | | | |
| | 10 ms WT | 1.77 ± 0.092 | 11/5 | WT vs R237W 10 ms | 0.745 | |
| | 10 ms R237W | 1.58 ± 0.09 | 10/6 | | | |
| | 20 ms WT | 1.88 ± 0.15 | 11/5 | WT vs R237W 20 ms | 0.874 | |
| | 20 ms R237W | 1.71 ± 0.096 | 10/6 | | | |
| | 50 ms WT | 1.78 ± 0.10 | 11/5 | WT vs R237W 50 ms | 0.399 | |
| | 50 ms R237W | 1.52 ± 0.082 | 10/6 | | | |
| | 100 ms WT | 1.71 ± 0.11 | 11/5 | WT vs R237W 100 ms | 0.358 | |
| | 100 ms R237W | 1.44 ± 0.099 | 10/6 | | | |
| | 200 ms WT | 1.41 ± 0.16 | 11/5 | WT vs R237W 200 ms | 0.677 | |
| | 200 ms R237W | 1.20 ± 0.024 | 10/6 | | | |
| | 500 ms WT | 1.17 ± 0.087 | 11/5 | WT vs R237W 500 ms | 0.999 | |
| | 500 ms R237W | 1.13 ± 0.069 | 10/6 | | | |
| **Fig 4e** | WT | | 34/20 | WT vs R237W Overall | 0.0181 | Two-way ANOVA |
| | R237W | | 39/23 | | | |
| | 25 μA WT | 68.6 ± 7.33 | 34/20 | WT vs R237W 25 μA | 0.132 | |
| | 25 μA R237W | 130.7 ± 18.12 | 39/23 | | | |
| | 50 μA WT | 143.3 ± 18.50 | 34/20 | WT vs R237W 50 μA | 0.0428 | |
| | 50 μA R237W | 237.7 ± 29.2 | 39/23 | | | |
| | 75 μA WT | 204.1 ± 25.6 | 34/20 | WT vs R237W 75 μA | 0.0049 | |
| | 75 μA R237W | 335.6 ± 37.3 | 39/23 | | | |
| | 100 μA WT | 249.4 ± 30.5 | 34/20 | WT vs R237W 100 μA | 0.0004 | |
| | 100 μA R237W | 499.0 ± 76.0 | 39/23 | | | |
| **Fig 4f** | WT | | 12/6 | WT vs R237W | 0.0043 | Two-way ANOVA |
| | R237W | | 13/6 | | | |
| **Fig 4g** | WT | | 12/6 | WT vs R237W | <0.0001 | |
| | R237W | | 13/6 | | | |
| **Fig 4h** | WT | 0.0548 ± 0.058 | 12/6 | WT vs R237W | 0.0045 | Mann-Whitney test |
| | R237W | 0.0328 ± 0.0041 | 13/6 | | | |
| **Fig 4i** | WT | 1.08 ± 0.12 | 12/6 | WT vs R237W | 0.739 | Unpaired t test |
| | R237W | 1.02 ± 0.15 | 13/6 | | | |
| **Fig 4j** | WT | 44.3 ± 3.8 | 12/6 | WT vs R237W | 0.0229 | |
| | R237W | 33.0 ± 2.7 | 13/6 | | | |
| Figure 6 | Group | Mean±SEM | n = # coverslips/ N = # of neuronal preparations | Comparison | P | Statistical test |

| | | | | | | |
|---|---|---|---|---|---|---|
| **Fig 6b** | DMSO | 0.348 ± 0.06 | 14/5 | DMSO vs 10 μM BMS | 0.0581 | One-way ANOVA with Dunnett's multiple comparison test |
| | 10 μM BMS | 0.145 ± 0.08 | 11/5 | | | |
| | 30 μM BMS | 0.130 ± 0.05 | 10/5 | DMSO vs 30 μM BMS | 0.0467 | |
| **Fig 6c** | DMSO | 0.508 ± 0.04 | 14/5 | DMSO vs 10 μM BMS | 0.202 | |
| | 10 μM BMS | 0.404 ± 0.04 | 11/5 | | | |
| | 30 μM BMS | 0.400 ± 0.06 | 10/5 | DMSO vs 30 μM BMS | 0.186 | |
| **Fig 6e** | +/+ DMSO | 0.109 ± 0.036 | 19/5 | +/+ DMSO vs +/+ BMS | 0.9876 | |
| | +/+ BMS | 0.126 ± 0.044 | 15/5 | | | |
| | R237W DMSO | 0.276 ± 0.034 | 16/5 | +/+ DMSO vs R237W DMSO | 0.0183 | |
| | R237W BMS | 0.033 ± 0.037 | 15/5 | +/+ DMSO vs R237W BMS | 0.4952 | |
| **Fig 6f** | +/+ DMSO | 0.410 ± 0.024 | 19/5 | +/+ DMSO vs +/+ BMS | 0.538 | |
| | +/+ BMS | 0.366 ± 0.030 | 15/5 | +/+ DMSO vs R237W DMSO | 0.925 | |
| | R237W DMSO | 0.428 ± 0.029 | 16/5 | +/+ DMSO vs R237W BMS | 0.089 | |
| | R237W BMS | 0.326 ± 0.023 | 15/5 | | | |

| Figure 7 | Group | Mean±SEM | n = # of coverslips/ N = # of neuronal preparations<br>n = # slices / N = # of animals | Comparison | P | Statistical test |
|---|---|---|---|---|---|---|
| **Fig 7a** | +/+ DMSO | | | +/+ DMSO vs +/+ BMS | 0.158 | Two-way ANOVA with Sidaks multiple comparison test |
| | +/+ BMS | | | | | |
| | 25 μA DMSO | 95.4 ± 13.5 | 13/6 | DMSO vs BMS 25 μA | >0.999 | |
| | 25 μA BMS | 102.5 ± 13.5 | 11/7 | | | |
| | 50 μA DMSO | 191.1 ± 35.5 | 13/6 | DMSO vs BMS 50 μA | 0.389 | |
| | 50 μA BMS | 288.5 ± 28.4 | 11/7 | | | |
| | 75 μA DMSO | 311.3 ± 48.9 | 13/6 | DMSO vs BMS 75 μA | 0.332 | |
| | 75 μA BMS | 414.6 ± 44.1 | 11/7 | | | |
| | 100 μA DMSO | 432.7 ± 68.2 | 13/6 | DMSO vs BMS 100 μA | 0.367 | |
| | 100 μA BMS | 532.4 ± 54.8 | 11/7 | | | |
| **Fig 7b** | +/+ DMSO | | | +/R237W DMSO vs +/R237W BMS | 0.0004 | |
| | +/+ BMS | | | | | |
| | 25 μA DMSO | 95.4 ± 13.5 | 13/6 | DMSO vs BMS 25 μA | 0.9546 | |
| | 25 μA BMS | 102.5 ± 13.5 | 13/7 | | | |
| | 50 μA DMSO | 191.1 ± 35.5 | 13/6 | DMSO vs BMS 50 μA | 0.7284 | |
| | 50 μA BMS | 288.5 ± 28.4 | 13/7 | | | |
| | 75 μA DMSO | 311.3 ± 48.9 | 13/6 | DMSO vs BMS 75 μA | 0.0410 | |
| | 75 μA BMS | 414.6 ± 44.1 | 13/7 | | | |
| | 100 μA DMSO | 432.7 ± 68.2 | 13/6 | DMSO vs BMS 100 μA | 0.0145 | |
| | 100 μA BMS | 532.4 ± 54.8 | 13/7 | | | |
| **Fig 7c** | +/+ DMSO | | 5/3 | +/+ DMSO vs +/+ BMS | 0.296 | Two-way repeated |

| | | | | | | |
|---|---|---|---|---|---|---|
| | +/+ BMS | | 6/4 | +/+ DMSO vs +/R237W DMSO | 0.002 | measures ANOVA |
| | +/R237W DMSO | | 5/3 | +/+ DMSO vs +/R237W BMS | 0.939 | |
| | +/R237W BMS | | 7/4 | +/R237W DMSO vs +/R237W BMS | 0.013 | |
| Figure 8 | Group | Mean±SEM | n = # animals | Comparison | P | Statistical test |
| **Fig 8b** | +/+ DMSO | 11.7 ± 4.8 | 14 | +/+ DMSO vs +/+ BMS | 0.586 | General linear model (repeated measures) with Bonferroni multiple comparison |
| | +/+ BMS | 15.6 ± 4.1 | 14 | +/+ DMSO vs +/R237W DMSO | 0.021 | |
| | +/R237W DMSO | 40.2 ± 10.5 | 14 | +/+ BMS vs +/R237W BMS | 0.314 | |
| | +/R237W BMS | 22.4 ± 5.2 | 14 | +/R237W DMSO vs +/R237W BMS | 0.019 | |
| **Fig 8c** | +/+ DMSO | 0.71 ± 0.45 | 14 | +/+ DMSO vs +/+ BMS | 0.393 | General linear model (repeated measures) with Bonferroni multiple comparison |
| | +/+ BMS | 2.21 ±0.96 | 14 | +/+ DMSO vs +/R237W DMSO | 0.016 | |
| | +/R237W DMSO | 7.14 ± 2.45 | 14 | +/+ BMS vs +/R237W BMS | 0.874 | |
| | +/R237W BMS | 2.43 ± 0.93 | 14 | +/R237W DMSO vs +/R237W BMS | 0.011 | |
| Figure S1 | Group | Mean±SEM | n = # of coverslips/ N = # of neuronal preparations | Comparison | P | Statistical test |
| **Fig S1** | Empty | 1.00 ± 0.089 | 16/4 | Empty vs WT | 0.0015 | One-way ANOVA with Dunnett's multiple comparison test |
| | WT | 2.663 ± 0.402 | 10/4 | | | |
| | K44A | 2.397 ± 0.353 | 11/4 | Empty vs K44A | 0.0065 | |
| | R237W | 2.759 ± 0.536 | 9/4 | Empty vs R237W | 0.0012 | |
| Figure S2 | Group | Mean±SEM | n = # of neuronal preparations or n = # of brains | Comparison | P | Statistical test |
| **Fig S2b** | +/+ Cell lysate | 1.000 ± 0.132 | 5 | +/+ vs +/R237W | 0.801 | Unpaired t test with Welch's correction |
| | +/R237W cell lysate | 0.918 ± 0.282 | 5 | | | |
| | +/+ 3 weeks | 1.000 ± 0.067 | 3 | +/+ vs +/R237W | 0.359 | |
| | +/R237W 3 weeks | 0.870 ± 0.112 | 3 | | | |
| | +/+ 6 weeks | 1.000 ± 0.115 | 3 | | 0.671 | |

| | | | | | | |
|---|---|---|---|---|---|---|
| | +/R237W 6 weeks | 0.955 ± 0.099 | 3 | +/+ vs +/R237W | | |
| **Fig S2d** | WT Syt1 | 1.00 ± 0.051 | 3 | +/+ vs +/R237W | 0.400 | Mann-Whitney test |
| | R237W Syt1 | 1.076 ± 0.041 | 3 | | | |
| | WT Amph1 | 1.00 ± 0.008 | 3 | +/+ vs +/R237W | 0.200 | |
| | R237W Amph1 | 1.111 ± 0.050 | 3 | | | |
| | WT C-Src | 1.00 ± 0.100 | 3 | +/+ vs +/R237W | 0.900 | |
| | R237W C-Src | 1.046 ± 0.057 | 3 | | | |
| | WT Endo | 1.00 ± 0.027 | 3 | +/+ vs +/R237W | 0.900 | |
| | R237W Endo | 1.044 ± 0.071 | 3 | | | |
| | WT Syd | 1.00 ± 0.040 | 3 | +/+ vs +/R237W | 0.700 | |
| | R237W Syd | 1.034 ± 0.037 | 3 | | | |
| | WT Eps15 | 1.00 ± 0.035 | 3 | +/+ vs +/R237W | 0.900 | |
| | R237W Eps15 | 1.029 ± 0.073 | 3 | | | |
| **Fig S2f** | WT Syt1 | 1.00 ± 0.142 | 3 | +/+ vs +/R237W | 0.700 | |
| | R237W Syt1 | 1.206 ± 0.222 | 3 | | | |
| | WT Amph1 | 1.00 ± 0.068 | 3 | +/+ vs +/R237W | 0.900 | |
| | R237W Amph1 | 1.103 ± 0.219 | 3 | | | |
| | WT C-Src | 1.00 ± 0.036 | 3 | +/+ vs +/R237W | 0.900 | |
| | R237W C-Src | 1.139 ± 0.203 | 3 | | | |
| | WT Endo | 1.00 ± 0.035 | 3 | +/+ vs +/R237W | 0.700 | |
| | R237W Endo | 1.184 ± 0.125 | 3 | | | |
| | WT Syd | 1.00 ± 0.002 | 3 | +/+ vs +/R237W | 0.700 | |
| | R237W Syd | 1.008 ± 0.095 | 3 | | | |
| | WT Eps15 | 1.00 ± 0.030 | 3 | +/+ vs +/R237W | 0.900 | |
| | R237W Eps15 | 1.057 ± 0.102 | 3 | | | |
| Figure S4 | Group | Mean±SEM | n = # of coverslips/ N = # of neuronal preparations | Comparison | P | Statistical test |
| **Fig S4b** | +/+ | 0.352 ± 0.019 | 11/3 | +/+ vs +/R237W | 0.839 | Unpaired t test |
| | +/R237W | 0.345 ± 0.028 | 10/3 | | | |
| **Fig S4d** | +/+ | 0.340 ± 0.016 | 17/3 | +/+ vs +/R237W | 0.919 | Mann-Whitney test |
| | +/R237W | 0.345 ± 0.021 | 16/3 | | | |
| **Fig S4e** | +/+ | 23072 ± 1556 | 8/4 | +/+ vs +/R237W | 0.974 | Unpaired t test |
| | +/R237W | 23145 ± 1571 | 7/4 | | | |
| Figure S5 | Group | Mean±SEM | n = # of coverslips/ N = # of neuronal preparations | Comparison | P | Statistical test |
| **Fig S5a** | DMSO | | 10/3 | DMSO vs 10 μM BMS | ns | Two-way ANOVA with Dunnett's multiple comparison test |
| | 10 μM BMS | | 8/3 | | | |
| | 30 μM BMS | | 6/3 | DMSO vs 30 μM BMS | <0.05 (100 s to 168 s) | |
| **Fig S5b** | DMSO | 0.323 ± 0.06 | 10/3 | DMSO vs 10 μM BMS | 0.424 | One-way ANOVA with Dunnett's multiple |
| | 10 μM BMS | 0.234 ± 0.05 | 8/3 | | | |
| | 30 μM BMS | 0.152 ± 0.04 | 6/3 | DMSO vs 30 μM BMS | 0.092 | |

| | | | | | | |
|---|---|---|---|---|---|---|
| **Fig S5c** | DMSO | 0.512 ± 0.04 | 10/3 | DMSO vs 10 μM BMS | 0.3894 | comparison test |
| | 10 μM BMS | 0.424 ± 0.02 | 8/3 | | | |
| | 30 μM BMS | 0.294 ± 0.05 | 6/3 | DMSO vs 30 μM BMS | 0.0217 | |

| Figure S6 | Group | Mean±SEM | n = # of coverslips/ N = # of neuronal preparations | Comparison | P | Statistical test |
|---|---|---|---|---|---|---|
| **Fig S6a(ii)** | DMSO | 0.334 ± 0.037 | 6/2 | DMSO vs 3 μM | 0.606 | One-way ANOVA with Dunnett's multiple comparison test |
| | 3 μM NS11021 | 0.386 ± 0.0235 | 7/2 | | | |
| | 10 μM NS11021 | 0.401 ± 0.030 | 8/2 | DMSO vs 10 μM | 0.356 | |
| | 30 μM NS11021 | 0.412 ± 0.040 | 6/2 | DMSO vs 30 μM | 0.337 | |
| **Fig S6a(iii)** | DMSO | 0.041 ± 0.022 | 6/2 | DMSO vs 3 μM | 0.987 | |
| | 3 μM NS11021 | 0.058 ± 0.044 | 7/2 | | | |
| | 10 μM NS11021 | 0.238 ± 0.032 | 8/2 | DMSO vs 10 μM | 0.011 | |
| | 30 μM NS11021 | 0.689 ± 0.069 | 6/2 | DMSO vs 30 μM | <0.0001 | |
| **Fig S6b(ii)** | DMSO | 0.393 ± 0.045 | 10/3 | DMSO vs 3 μM | 0.902 | |
| | 3 μM Retigabine | 0.424 ± 0.038 | 13/3 | | | |
| | 10 μM Retigabine | 0.428 ± 0.026 | 14/3 | DMSO vs 10 μM | 0.868 | |
| | 30 μM Retigabine | 0.456 ± 0.051 | 10/3 | DMSO vs 30 μM | 0.597 | |
| **Fig S6b(iii)** | DMSO | 0.084 ± 0.046 | 10/3 | DMSO vs 3 μM | >0.999 | |
| | 3 μM Retigabine | 0.084 ± 0.038 | 13/3 | | | |
| | 10 μM Retigabine | 0.072 ± 0.033 | 14/3 | DMSO vs 10 μM | 0.991 | |
| | 30 μM Retigabine | 0.216 ± 0.031 | 10/3 | DMSO vs 30 μM | 0.054 | |
| **Fig S6c(ii)** | DMSO | 0.366 ± 0.037 | 9/4 | DMSO vs 3 μM | 0.471 | |
| | 3 μM BMS191011 | 0.306 ± 0.044 | 10/4 | | | |
| | 10 μM BMS191011 | 0.310 ± 0.031 | 10/4 | DMSO vs 10 μM | 0.513 | |
| | 30 μM BMS191011 | 0.276 ± 0.020 | 12/4 | DMSO vs 30 μM | 0.145 | |
| **Fig S6c(iii)** | DMSO | 0.074 ± 0.074 | 9/4 | DMSO vs 3 μM | 0.821 | |
| | 3 μM BMS191011 | 0.009 ± 0.056 | 10/4 | | | |
| | 10 μM BMS191011 | 0.026 ± 0.062 | 10/4 | DMSO vs 10 μM | 0.917 | |
| | 30 μM BMS191011 | 0.067 ± 0.054 | 12/4 | DMSO vs 30 μM | 0.999 | |
| **Fig S6d(ii)** | DMSO | 0.481 ± 0.030 | 15/3 | DMSO vs 10 μM | 0.003 | Unpaired t test |
| | 10 μM Paxilline | 0.376 ± 0.012 | 15/3 | | | |
| | DMSO | 0.115 ± 0.038 | 15/3 | | 0.427 | |

| | | | | | | |
|---|---|---|---|---|---|---|
| **Fig S6d(iii)** | 10 $\mu$M Paxilline | 0.072 ± 0.037 | 15/3 | DMSO vs 10 $\mu$M | | |
| **Fig S6e(ii)** | DMSO | 0.456 ± 0.031 | 13/3 | DMSO vs 6 $\mu$M | 0.361 | |
| | 6 $\mu$M XE-991 | 0.406 ± 0.039 | 13/3 | | | |
| **Fig S6e(iii)** | DMSO | 0.123 ± 0.056 | 13/3 | DMSO vs 6 $\mu$M | 0.152 | Mann-Whitney test |
| | 6 $\mu$M XE-991 | 0.022 ± 0.047 | 13/3 | | | |

| Figure S7 | Group | Mean±SEM | n = # animals | Comparison | P | Statistical test |
|---|---|---|---|---|---|---|
| **Fig S7a** | +/+ | 25.29 ± 5.61 | 14 | +/+ vs +/R237W | 0.0066 | Unpaired t test with Welch's correction |
| | +/R237W | 59.71 ± 9.92 | 14 | | | |
| **Fig S7b** | +/+ | 3.72 ± 1.25 | 14 | +/+ vs +/R237W | 0.0238 | |
| | +/R237W | 10.14 ± 2.31 | 14 | | | |
| **Fig S7c** | +/+ DMSO | 136.0 ± 5.1 | 14 | N/A | 0.198 | General linear model (repeated measures) |
| | +/+ BMS | 138.2 ± 6.3 | 14 | | | |
| | +/R237W DMSO | 161.1 ± 8.2 | 14 | | | |
| | +/R237W BMS | 151.4 ± 5.7 | 14 | | | |
| **Fig S7d** | +/+ DMSO | 36.2 ± 1.9 | 14 | N/A | 0.550 | General linear model (repeated measures) |
| | +/+ BMS | 34.2 ± 1.4 | 14 | | | |
| | +/R237W DMSO | 37.0 ± 1.0 | 14 | | | |
| | +/R237W BMS | 36.5 ± 1.5 | 14 | | | |

**Extended Data Table 4** – *Collated table of experimental n, P values and statistical tests*.

**Extended Data Movie 1** – *Example of myoclonic jumping in the $Dnm1^{+/R237W}$ mouse*. Movie displaying the myoclonic jumping phenotype. Myoclonic jumping can be observed at 18, 32, 35, 113,114, 119, 123, 124, 126, 129, 131, 132, 134, 135, 136, 147, and 175 s with bursts occurring between 113-114, 123-126 and 131-136 s.
